# Likelihood-ratio test statistic for the finite-sample case in nonlinear ordinary differential equation models

**DOI:** 10.1101/2023.03.25.534223

**Authors:** Christian Tönsing, Bernhard Steiert, Jens Timmer, Clemens Kreutz

## Abstract

Likelihood ratios are frequently utilized as basis for statistical tests, for model selection criteria and for assessing parameter and prediction uncertainties, e.g. using the profile likelihood. However, translating these likelihood ratios into p-values or confidence intervals requires the exact form of the test statistic’s distribution. The lack of knowledge about this distribution for nonlinear ordinary differential equation (ODE) models requires an approximation which assumes the so-called asymptotic setting, i.e. a sufficiently large amount of data. Since the amount of data from quantitative molecular biology is typically limited in applications, this finite-sample case regularly occurs for mechanistic models of dynamical systems, e.g. biochemical reaction networks or infectious disease models. Thus, it is unclear whether the standard approach of using statistical thresholds derived for the asymptotic large-sample setting in realistic applications results in valid conclusions. In this study, empirical likelihood ratios for parameters from 19 published nonlinear ODE benchmark models are investigated using a resampling approach for the original data designs. Their distributions are compared to the asymptotic approximation and statistical thresholds are checked for conservativeness. It turns out, that corrections of the likelihood ratios in such finite-sample applications are required in order to avoid anti-conservative results.

**Author summary:** Statistical methods based on the likelihood ratio are ubiquitous in mathematical modelling in systems biology. For example confidence intervals of estimated parameters rely on the statistical properties of the likelihood-ratio test. However, it is often overlooked that these intervals sizes rely on assumptions on the amounts of data, which are regularly violated in typical applications in systems biology. By checking the appropriateness of these assumptions in models from the literature, this study shows that in a surprisingly large fraction confidence intervals might be too small. Using a geometric interpretation of parameter estimation in the so-called data space, it is motivated why these issues appear and how they depend on the identifiability of the model parameters. In order to avoid such problematic situations, this work makes suggestions on how to adapt the statistical threshold values for likelihood-ratio test. By this, it can be assured that valid statistical conclusions are drawn from the analysis, also in situations where only smaller data sets are available. Such corrections yield for example more conservative confidence interval sizes and thus decrease a potential underestimation of the parameter uncertainty.

## Introduction

An essential characteristic of systems biology is measurement-driven learning about a biological process of interest [1, 2]. A typical approach is translating existing knowledge about biochemical interactions in living cells into mathematical models. Then, experimental data is utilized to check whether the current understanding of a system as represented in the model is in agreement with observations. In the case of concordance, model parameters can be estimated, the system’s response can be studied by computational approaches, and its behaviour can be predicted by the model. In the case of mismatch between data and mathematical model, new interactions and hypotheses might be suggested and tested. For both scenarios, a proper and efficient statistical analysis is required to assess significance and prevent invalid conclusions.

For traditional comparisons which are not based on dynamic models, a large number of well-established statistical tests is available for assessing significance. Some of these tests like the *t-test* or *analysis of variance* (ANOVA) provide valid results for any sample size. Other tests like for example *Pearson’s chi-squared test* for independence are derived under the so-called asymptotic assumption, which implies that they are only valid for a large sample size and only provide approximative results for the finite-sample case when only limited amounts of data are available [3].

The *likelihood* is the probability of a data set given some hypothesis which can be applied to assess agreement of data and model for any kind of noise distribution. The so-called *likelihood principle* states that all information contained in the data concerning two hypotheses is comprised in the likelihood ratio of those hypotheses [4]. Moreover, the *Neyman-Pearson lemma* guarantees that if the assumptions about a null model are valid, tests based on likelhood ratios have maximal power compared to competing test statistics [5]. This is the reason, why many traditional tests can be derived by considering likelihood ratios or can be interpreted in terms of a likelihood-ratio test. This theoretical statement, however, does not provide generally applicable formulas for translating likelihood ratios into p-values or thresholds of the test statistic.

A general statement is only available for the asymptotic, large-sample setting which is usually applied but not fulfilled in systems biology [6]. For typical modelling applications in systems biology, e.g. for ordinary differential equation (ODE) models, there are no traditional statistical tests available which are valid in the finite-sample case. Moreover, models can also be ranked based on their agreement with the data using other criteria like the Akaike information criterion (AIC) or the Bayesian information criterion (BIC) and variants thereof are available [7].

The likelihood-ratio test is applied for discriminating between nested models. Moreover, a number of statistical methods related to likelihood ratios are utilized, for example confidence intervals for parameters can be calculated based on the profile likelihood which is a continuous representation of the likelihood-ratio statistic. This approach can also be used for investigation of parameter identifiability [8], observability analysis [6] as well as for model reduction [9, 10]. To apply the likelihood-ratio test in such cases, typically the asymptotic theory for the test statistic is utilized for the finite-sample case, but without checking the appropriateness of the description.

In this work, the appropriateness of such asymptotic assumptions for the distribution of the likelihood-ratio test is checked using 19 nonlinear ODE benchmark models and realistic data designs from the literature. Using a parametric bootstrapping procedure, artificial, yet realistic data realisations are generated and empirical likelihood ratios for practical application scenarios are simulated. The main goal of this work is to evaluate how frequently deviations from the asymptotic large-sample assumptions occur in these situations.

We observe differences in the resulting distributions to the expected asymptotic theory that might be relevant for the drawn biological conclusions by the dynamic models. Using a geometrical interpretation of parameter estimation in such models in the data space, we present an explanation how certain model structures cause such deviations. To cure the issue in realistic scenarios, corrections of the likelihood ratios are suggested which avoid anti-conservative results of the statistical methods. Moreover, potential adaptations for alternative statistical thresholds are given and an approach for the re-interpretation of existing studies, which use the asymptotic threshold, is presented.

## Methods

### Parameter estimation

Mechanistic models of biological systems can be formulated by ordinary differential equations (ODEs)

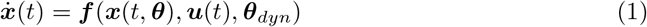

that describe the dynamics of the models states ***x*** depending on the model structure and specifically on the dynamic parameters **θ**_*dyn*_, e.g. reaction rate constants. External effects can be incorporated by an input function ***u***(*t*). The initial value problem can be solved numerically and yields solutions ***x***(*t*) that are non-linear with respect to the parameters. Since the specific parameter values of the systems are unknown in applications, they need to be estimated from experimental data. Also initial values of the ODE system ***x***(*t* = 0) = ***θ***_init_ can be treated as unknown parameters and can be estimated simultaneously [11].

Typically, quantification of biological counterparts corresponds to sums or fractions of the model states ***x***(*t*) and involves indirect measurements of the observed entities such as for instance relative data. Thus, the experimental data is compared to the model trajectories by

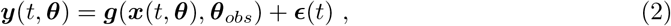

i.e. by observation functions ***g*** that incorporate observational parameters ***θ***_obs_ such as scalings and offsets of the quantified signal. Typically Gaussian noise 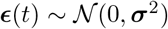 with a fixed standard deviation *σ* is assumed here, but in general also more complex error models for ***σ*** = ***σ***(***x***(*t*), ***θ**_σ_*) with error model parameters ***θ**_σ_* might be considered which yield the desired noise distribution. In applications, systems are often only partially observed as the quality and amount of the data is limited and only sparse data sets are available. The experimental design 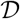 summarizes the measured observables, experimental conditions, temporal sampling points as well as potential replicate measurements [12].

In systems biology related models the values of the parameters

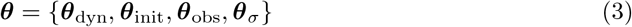

often cannot be directly inferred but need to be estimated based on a mathematical model and experimental data [13]. The maximum likelihood estimate (MLE)

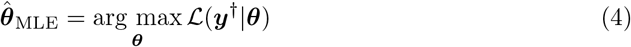

is the most efficient and asymptotically unbiased estimator. It is the parameter vector that maximizes a likelihood function given the experimental data ***y***^†^ or data realisation of the experimental design 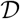 in a simulation setting [14]. For parameter estimation, equivalently the so-called log-likelihood function 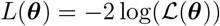 can be minimized, which is often numerically advantageous [11]. The log-likelihood function

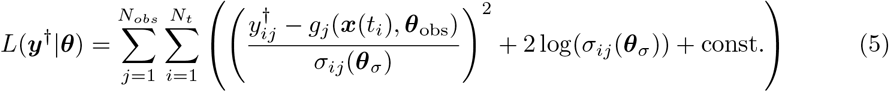

quantifies the discrepancy between model output and data 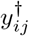 at the *i*-th timepoint and *j*-th observable. In the special case of an error model that assumes Gaussian noise, this is equivalent to the method of weighted least squares [15, 16].

### Likelihood-ratio test

For testing hypotheses in parametric models, the *likelihood-ratio test* can be used [17]. It can be utilized for example to compare two alternative model structures based on their likelihood functions, as they comprise all available information about the model and data with respect to the parameters.

Consider a full model 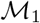 with parameters ***θ***_1_ ∈ Θ_1_ and an alternative model 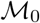 that is a nested simplification thereof with parameters ***θ***_0_ ∈ Θ_0_, i.e. Θ_0_ ⊂ Θ_1_. The null hypothesis *H*_0_ is formulated such that Θ_0_ in 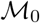 is a valid simplification of Θ_1_ in 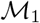. If *H*_0_ is rejected, it can be concluded that the simplification is not valid. The respective likelihood ratio

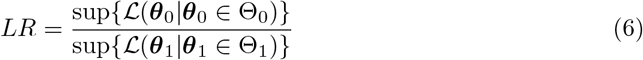

yields the test statistic

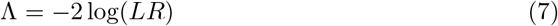

with 0 < *LR* ≤ 1, such that 0 < Λ < ∞ [14]. In the following, mentioning the supremum function is omitted for readability as always the *optimized* likelihood function is assumed. That is the likelihood function 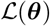 at the MLE 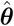, i.e. where its value is maximal, or minimal in terms of the log-likelihood 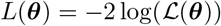. Using the latter expression, the test statistic

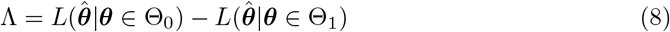

reduces to a simple difference of the log-likelihoods *L*.

Depending on the value of the test statistic, the null hypothesis *H*_0_ should be rejected if the likelihood ratio Λ exceeds a critical threshold 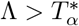. However, except from special cases, the distribution of the likelihood ratios and its quantile function cannot be derived theoretically. Thus, in applications the exact relationship between critical value and the significance level *α* is unknown. As a consequence, translating likelihood ratios into p-values or determining a critical value *T_α_* of the test statistic remains problematic for realistic applications.

Under mild assumptions on the regularity, this issue can be circumvented by using *Wilks’ theorem* to approximate the distribution of the likelihood-ratio test statistic in the asymptotic case. For this, assume that model 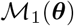 with ***θ*** ∈ Θ_1_ has *r* free parameters, while the nested model 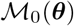 with ***θ*** ∈ Θ_0_ has *s* free parameters with *s* < *r*, that the nested simplification Θ_0_ ⊂ Θ_1_ is specified appropriately, that all parameters are identifiable and that the true parameters are not at the boundary of the parameter space [18]. Then, for a sufficiently large number of data points, the test statistic

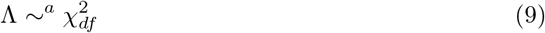

asymptotically converges to a chi-square distribution 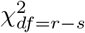 with *r* – *s* degrees of freedom equal to the difference of free model parameters *r* – *s*, independently from the true parameter value and independently from the underlying likelihood functions.

By this, the probability distribution of the test statistic Λ in the asymptotic case is available from the density function of the *χ*^2^-distribution *ρ*_χ^2^_ (*x, df*). Given a significance level *α* or confidence level 1 – *α*, thresholds 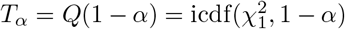 of the test statistic can be determined using the quantile function or inverse cumulative distribution function of the *χ*^2^-distribution in the large sample case.

#### Profile likelihood

Confidence intervals for point estimates are traditionally calculated from the Fisher information matrix (FIM), which is derived from second-order derivatives of the likelihood at the MLE and thus represents the local curvature of the likelihood [19, 20]. This yields exact results for linear models but does not account for non-zero higher-order derivatives of the likelihood with respect to the parameters. In contrast, the *profile likelihood* approach represents a generalisation for nonlinear systems. To calculate the profile likelihood

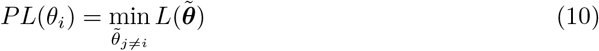

for one parameter of interest *θ_i_* that is scanned along its axis around the MLE while re-optimizing all remaining parameters *θ*_*i*≠*j*_. Since this approach directly evaluates the likelihood function, it enables the calculation of reliable confidence intervals of the estimated parameters also for nonlinear models [8, 21–23].

In analogy to FIM-based standard errors motivated from the analysis of linear models, the uncertainty of parameter estimates of nonlinear ODE models can be assessed by the profile likelihood-based confidence interval

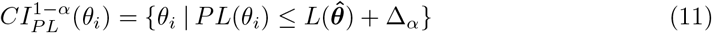

for a confidence level 1 – *α*. It represents the set of acceptable parameter values, characterized by a profile likelihood value *PL*(*θ_i_*) below the threshold Δ_*α*_ compared to the log-likelihood value 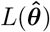 at the MLE 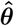.

This confidence interval can be interpreted as a continuous likelihood-ratio statistic

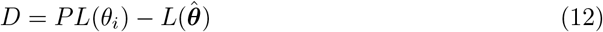

for which the null hypothesis cannot be rejected if *D* < Δ_*α*_, following the discussed case with a simplification of nested models with one fixed parameter. Using *Wilks’ theorem*, the critical value of the test statistic *T_α_* can be analogously defined. In this line, the threshold Δ_*α*_ is calculated under the asymptotic assumption *N_data_* → ∞ from the (1 – *α*)-quantile, i.e. 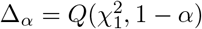 using the 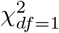-distribution with one degree of freedom.

In addition to the definition of confidence intervals, the profile likelihood approach also allows for a data-based identifiability analysis of the estimated parameters. Based on the shape of the likelihood profile a classification scheme is available, that again depends on the the statistical threshold in terms of the log-likelihood [8, 24]. A parameter is considered as *practically identifiable* if its likelihood profile exceeds the statistical threshold given by 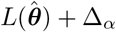 for values of *θ_i_* smaller *and* larger than the MLE 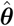, i.e. yielding finite profile likelihood-based confidence intervals, as depicted in Fig. 1 A. If the amount or quality of the available data is not sufficient to indicate a confidence bound in both directions of *θ_i_*, the parameter is termed as *practically non-identifiable*. In this case, a unique point estimate may be obtained, but the profile does not exceed the threshold *Δ_α_* in at least one direction and thus does not have a confidence interval with finite bounds, c.f. Fig. 1 B. A constant profile likelihood for the whole parameter domain as shown in Fig. 1 C indicates a *structurally non-identifiable* parameter. Such a case arrises if the estimated parameters are not unique for the given data set, for example if the model structure contains a redundant parametrization which cannot be resolved by additional data.

**Fig 1.**
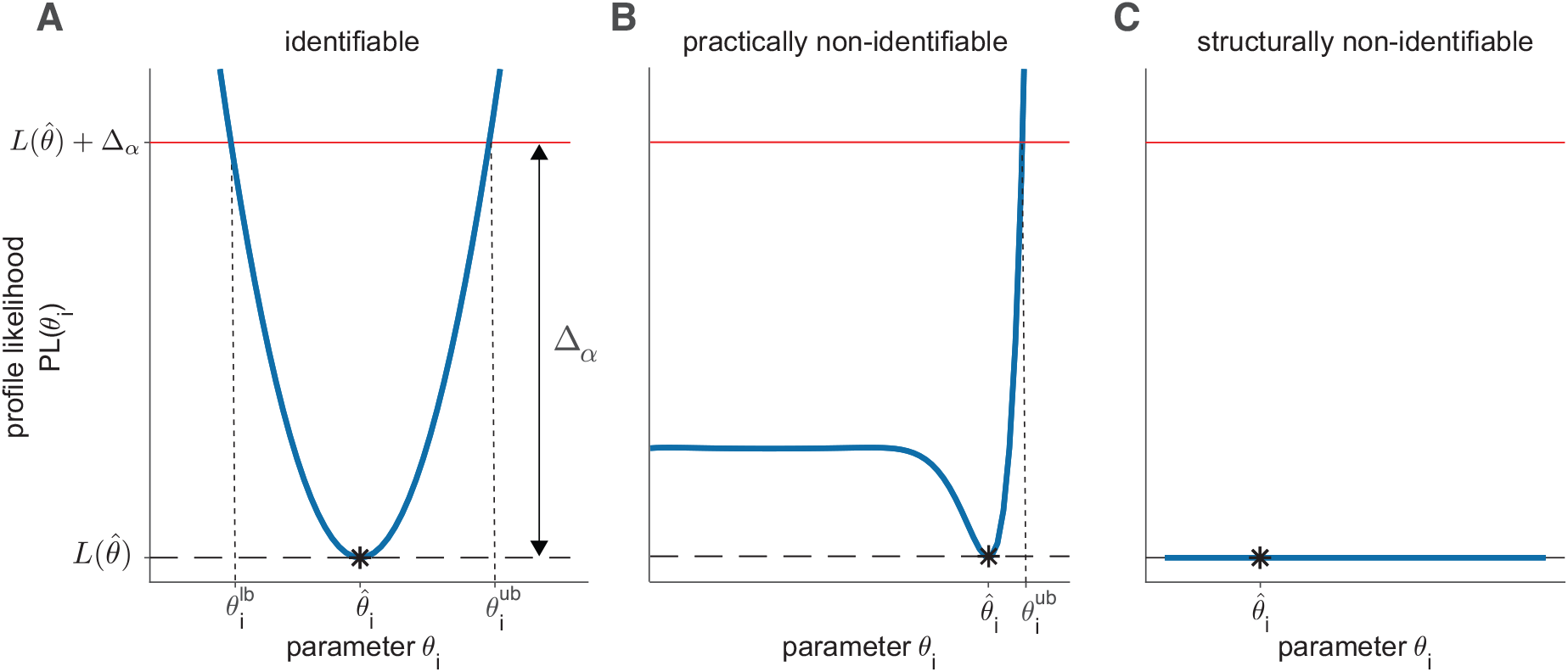
Profile likelihood for identifiability analysis. (A) Illustrative profile likelihood as blue solid line for an identifiable parameter. For analyses of the identifiability status, the intersection with statistical threshold value (red line) at 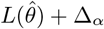 is crucial. Furthermore, profile likelihood-based confidence intervals can be constructed from the projections of the intersections on the parameter axis, defining the lower *θ_lb_* and upper bound *θ_ub_* of the confidence interval, if these intersections exist. The MLE 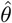 is indicated by the black asterisk. (B) Profile likelihood of a practically non-identifiable parameter that is open to the left resulting in an unbounded confidence interval. (C) Likelihood profile of a structural non-identifiability indicated by a flat line with unbounded confidence interval in both directions.

### Corrections of likelihood ratios for finite-data samples

In realistic modeling applications, only a limited number of data points are available, i.e. in contrast to the assumed asymptotic case, the finite-sample case typically occurs. Thus, the validity of the approximation of the likelihood-ratio test statistic with the 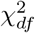-distribution from Eq. (9) via Wilks’ theorem cannot be guaranteed in general.

#### Bartlett correction

An improved approximation of such a test statistic by a *χ*^2^-distribution has been proposed by Bartlett [25]. It relies on a problem-specific Barlett correction factor *C* = 1/(1 + *df/q*), where in the original formulation *df* is the difference between the dimensions of the parameter spaces of the model under the alternative hypothesis and the null hypothesis and *q* is a constant which needs to be determined for the individual application. It has been shown that the modified likelihood ratio statistic

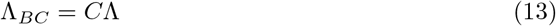

is closer to the 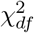-distribution in applications than the unmodified statistic, i.e. the first three cumulants agree up to an error of 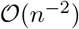 [26]. Depending on the actual setting and model formulation, the appropriate Bartlett correction factor is difficult to obtain or may be impossible to compute in realistic applications [17]. The idea of the *Bartlett correction* was later generalized by Lawley [27] with a generalized expression for the correction factor *C* as a function of the moments of the log-likelihood derivatives. The analytical derivation of Lawley’s approach, however is cumbersome, as it requires joint cumulants of log-likelihood derivatives up to the fourth order. While there are adaptions for certain model classes that allow a more efficient calculation of the respective Bartlett correction factor [28–31], no methods are available for nonlinear ODE models.

#### Chebyshev’s and Cantelli’s inequality

Critical values of the test statistic are essential as statistical thresholds for the discussed likelihood-ratio test related methods. In addition to their derivation based on Wilk’s theorem and calculation from the *χ*^2^-distribution, we propose an alternative and more general attempt of defining such thresholds using Chebyshev’s and Cantelli’s inequality.

For a simple illustration, let’s first consider a random variable 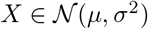 generated from a normal distribution with mean *μ* and standard deviation *σ*. Using the so-called *empirical rule*, for example for the 3*σ* neighborhood, the probability *P*(|*X* – *μ* | < 3*σ*) is known to be 99.7%. A popular implication from this rule is that almost all of the variables of a respective data sample are contained in the 3*σ* interval, if the numbers are normally distributed [32]. Moreover, it is approximately valid if the underlying distribution is at least bell-shaped with mean *μ* and standard deviation *σ*. Assuming that the distribution of a two-tailed test statistic would follow such a (approximative) normal distribution, statistical threshold could be constructed from the empirical rule according to a significance level *α* = 1 – *P*. For such a two-tailed test statistic a lower and upper statistical threshold 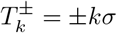 could be used as lower and upper critical values for the two rejection regions at the outer parts of the distribution’s tails.

Under much weaker assumptions on the distribution, *Chebyshev’s inequality* can be utilized[33]. If the empirical mean *μ* and standard deviation *σ* of the sample are finite and known, Chebyshev’s inequality guarantees

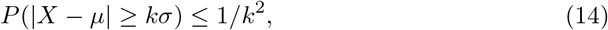

i.e. in average *at least* (1 – 1/*k*^2^) × 100% of random numbers X drawn from the distribution to be contained in the interval [*μ* – *kσ*, *μ* + *kσ*]. This is completely independent from the underlying probability distribution and is thus also suitable even in cases when the form of the probability distribution is unknown.

It can be concluded that a realisation of a random number X of any kind of distribution lies beyond *k* = 4.47 standard deviations of its mean in *a maximum of* 5% of the cases, i.e. with *α* ≤ 5 %. Again, for the rejection region of an hypothesis test an upper bound 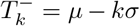 on the left tail of the distribution can be defined, as well as a lower bound 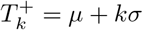 for the rejection region on the right tail. Chebyshev’s inequality can be also applied to a positive one-tailed distribution, if 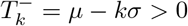, which is valid for the 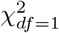-distribution for 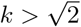.

For random variables drawn from single-tailed distributions, a one-sided threshold can be similarly derived from *Cantelli’s inequality* [34]. For a realisation of random numbers *X* in a sample with empirical mean *μ* and standard deviation *σ*, the probability that a single random number exceeds the sample’s mean by any real number *c* > 0 is

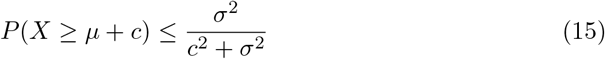

regardless of the underlying probability distribution. Motivated from the conclusion of Wilk’s theorem that the likelihood ratios should be at least asymptotically 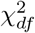-distributed, Cantelli’s inequality can be used to derive a value for the statistical threshold in the finite-sample case, i.e. when a certain deviation from the asymptotic distribution is expected. For this, assume an in detail unknown but 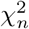-like distribution with mean *μ* = *n* and standard deviation 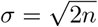. It then follows from Eq. (15) that a realisation of a random number X from such a distribution exceeds a critical value of *T_ξ_* = *μ* + *c* with probability *α* ≤ 2*n*/(*c*^2^ + 2*n*). The same rationale then applies when assuming an in detail unknown test statistic, which has 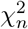-like mean *μ* = *n* and standard deviation 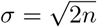. The critical value threshold *T_ξ_* of this test statistic can then be calculated from Cantelli’s inequality for a significance level of maximal *α* ≤ 2*n*/(*c*^2^ + 2*n*). In conclusion and for *n* = 1 degrees of freedom, this yields a critical value *T_ξ_* = 1 + 6.16 = 7.16 for the statistical threshold for a maximal 5% significance level under the assumption of the in detail unknown, but 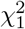-like test statistic. Of note: the threshold of a 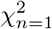-distribution for a significance level *α* = 0.05 would be 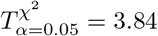

## Approach

### Realistic data simulation approach

To check the validity of the asymptotic *χ*^2^-distribution of the likelihood ratio from Wilks’ theorem as well as the appropriateness of alternative threshold formulations, empirical likelihood ratios form realistic model examples are employed. For this, in total 19 nonlinear ODE benchmark models of different biological systems and with various model properties are used. Additional data sets are simulated according to the model’s original experimental design using parametric bootstrapping. The thereof resulting empirical likelihood ratios for each parameter are used for comparisons to asymptotic theory and a classification scheme based on probability-probability plots is developed to assess the conservativeness of the likelihood ratio statistic.

#### Parametric bootstrapping

Bootstrapping is a frequently applied method to estimate the sampling distribution of an arbitrary statistic [35]. For parametric bootstrap, the MLE 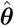 is assumed as underlying truth and used for generation of realisations of artificial data samples from the distribution of the measurement errors. For the nonlinear ODE models analyzed here, model trajectories are simulated from the observables according to the original experimental design 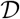 using the MLE of the original data set. For each model, *N_real_* = 500 realisations of artificial data sets are generated by adding noise to the simulations according to the original error model.

Then, according to the parametric bootstrapping proceedure the model parameters are re-estimated for a data realization 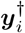, yielding the value of the log-likelihood function termed as 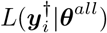. To check the distribution of the likelihood ratios for the earlier discussed case of nested models with one fixed parameter, the log-likelihood value termed as 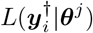 is determined for the respective data realization 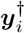 and for all parameters *θ_j_*. This is done by fixing each parameter of interest *θ_j_* to the value obtained from the fit to the original data, while re-fitting the model by re-optimizing all remaining parameters.

Since the parameter of interest is fixed to its true value 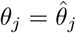, i.e. to the value which generates the model trajectories from which the artificial data sets are sampled, the model with *L*(***θ***|***θ*** ∈ Θ_0_) with nesting 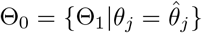 is a valid simplification of the full model with likelihood function *L*(***θ***|***θ*** ∈ Θ_1_). Thus, the likelihood ratio or the difference in terms of log-likelihood *L* of both fits, i.e. the fit of all parameters and the fit with fixed *j*-th parameter *θ_j_* yields

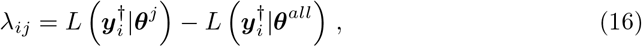

for each data realization 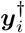. According to *Wilks’ theorem*, the distribution of the empirical likelihood ratios

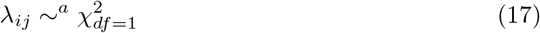

should asymptotically hold.

The described bootstrapping procedure as summarized in pseudocode in S1 Fig is repeated for each of the *N_real_* = 500 data realisations and for each estimated parameter *θ_j_* of the *m*-th model, yielding the empirical realization of the likelihood ratio *λ_ijm_* for the *i*-th data realization. By calculating the empirical cumulative density function (ECDF) of *λ_ijm_* over all simulated data samples, the empirical distribution can be related to the theoretically expected cumulative density function (CDF), for example to the expected 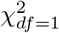-distribution in order to check the appropriateness of the asymptotic approximation. For this, empirical thresholds of *λ* for different significance levels *α* can be compared to the expected quantiles 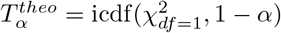.

### Classification of empirical likelihood ratios

In order to draw practically relevant conclusions about the distribution of the empirical likelihood ratios from the parametric bootstrapping procedure, the empirical cumulative density function (ECDF) is investigated. A classification scheme is carried out in order to compare the empirical distribution to the theoretically expected distribution. By this, we check the appropriateness of Wilks’ theorem for thresholds of the likelihood-ratio test statistic and related statistical methods.

#### Probability-probability plots (pp-plots)

To compare the ECDF of the empirical likelihood ratio samples against the theoretical CDF of the asymptotically expected reference distribution, *probability-probability-plots (pp-plots)* are utilized [36]. If the sample is distributed according to the theoretically expected distribution, the identical cumulative probabilities for the respective quantiles in both the theoretical CDF and in the ECDF will be observed. For visual inspection, the pp-plot is constructed by plotting the ECDF of the likelihood ratio sample, i.e. the cumulative probability values on the y-axis against the cumulative probability values of the theoretical CDF on the x-axis of the pp-plot, cf. Fig. 2 A. A data sample distributed accordingly to the theoretical CDF would lie on the diagonal, as shown by the red line in Fig. 2 A. This outcome is expected for the asymptotic case when the empirical likelihood ratio sample is indeed distributed as 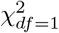.

**Fig 2.**
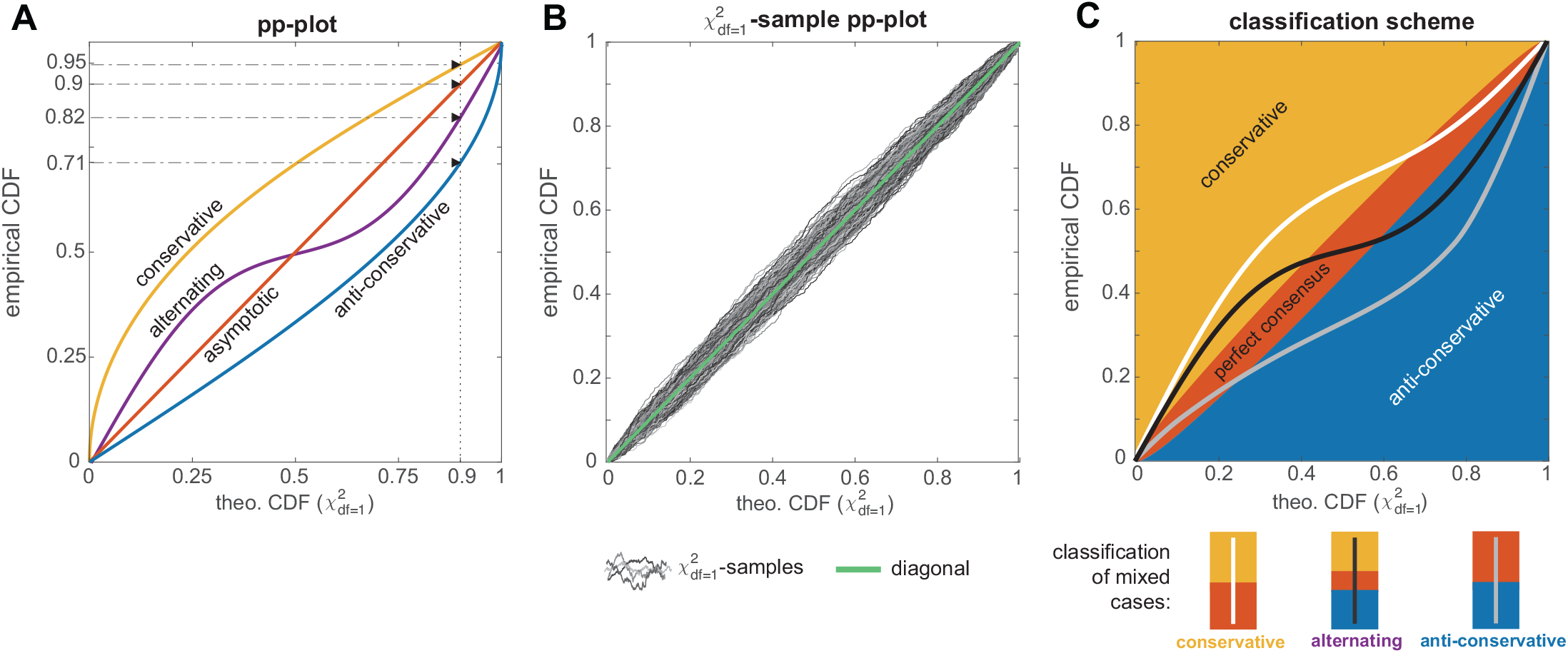
Empirical likelihood ratio classification. A: Probability-probability plot (pp-plot) with four possible graph scenarios. B: 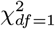-samples show a noticeable deviation from the diagonal in the pp-plot. The pp-plot in panel B shows graphs from 1000 samples each with *N_real_* = 500 random numbers drawn from a 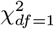-distribution. C: Classifications plot with classification regions and classification scheme for mixed cases: the white graph is classified as *conservative*, although its upper part lies in the perfect consensus area. Likewise, the light gray graph is identified with the *anti-conservative* case, while the black graph represents an *alternating* case which populates all three areas.

If however, the cumulative probability of the ECDF is larger than the expected theoretical CDF, the pp-plot graph lies in the upper left triangle. This case is termed *conservative*, since the ECDF implies a smaller empirical threshold value 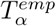 for the significance level *α*, compared the expected threshold 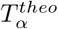 from the theoretical CDF. That is, the same likelihood ratio corresponds to a larger cumulative probability in the ECDF than for the theoretical CDF. In other words, the empirical sample contains more smaller values of the likelihood ratios than expected, i.e. more than the expected 1 – *α*% of the likelihood ratio values are smaller than the expected 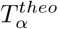 of the reference distribution. The consequence of applying such a non-adequate threshold 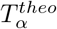 derived from the asymptotic theory to this sample would be for example a too conservative confidence interval. A respective profile likelihood-based confidence interval with a threshold 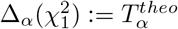 would be too large and the true value of a parameter would lie more often than in 1 – *α*% of the cases within the confidence interval. The pp-plot graph of such a conservative case is shown in yellow in Fig. 2 A. It indicates that the ECDF has a larger cumulative probability value compared to the value of the theoretical CDF at the chosen confidence level, cf. the example for a confidence level 1 – *α* = 0.9 in Fig. 2 A. For the shown example, *more* than 90% of the data sample are in fact smaller than the expected 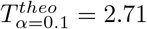.

In contrast, the opposite case of a too small threshold is termed *anti-conservative*and is identified by a pp-plot graph lying entirely in the lower right triangle, c.f. blue line in Fig. 2 A. Here, a profile likelihood-based confidence interval with 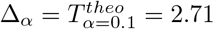 would be too small, since in this example only 71% of the likelihood ratios are smaller than *T^theo^*, in contrast to the expected 90%.

Taking into account the course of the whole pp-plot graph, also a mixture of both cases can occur, as shown by the purple line in Fig. 2 A. In this *alternating* case, the graph lies for example in the conservative triangle for small values of the CDF, but crosses the diagonal in the middle of the cumulative probability domain of the CDF and remains below the diagonal for larger *p*-values in the anti-conservative regime.

#### Classification of realistic samples

For the further analysis, a classification scheme for the pp-plots of the empirical likelihood ratios according to the discussed cases is implemented. This is necessary since realistic samples are less smooth than the graphs depicted in Fig. 2A and even realisations from a 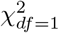-distributed random variable show a noticeable deviation from the expected straight diagonal in the pp-plot (light green line in Fig. 2B). For illustration, 1000 random samples each containing *N_real_* = 500 i.i.d. random variables drawn from the expected 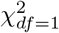-distribution are generated, resulting in pp-plot graphs depicted in shades of gray in Fig. 2 B. In order to be able to identify empirical likelihood ratios which in fact correspond to the asymptotic theory, but show a similar deviation from the diagonal, a tolerance region around the diagonal is constructed for the classification scheme. This area is constructed from fitting a higher-order polynomial around the 3*σ*-deviation of the 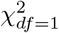 samples around the diagonal This polynomial is bounded at *p* = 0 and 1 and shows the largest deviation from the baseline at *p* = 0.5. The resulting tolerance band contains approximately 95% of the 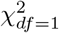-sample graphs. In the context of the bootstrapping procedure for empirical likelihood ratios, a graph which remains entirely within this area of the pp-plot is termed as a *perfect consensus*with the asymptotic theory and is classified accordingly.

Graphs lying entirely above or below the red *perfect consensus*-area are identified either with the *conservative* case (yellow) or *anti-conservative* case (blue), respectively. Despite these clear cases, also mixed cases needs to be considered. Often, the conservative or anti-conservative graphs do not lie exclusively in their respective areas, but their tails overlap with the perfect consensus-area. Such a pp-plot graph with a deviation from perfect consensus-area in only one direction is assigned to the respective class. For example the white graph in Fig. 2C is be assigned to the conservative case, even if parts of it lie in the perfect consensus-area, but not in the anti-conservative area. Likewise, the gray graph is identified as an anti-conservative case.

Contrarily, graphs that have parts in both, the conservative and the anti-conservative part are assigned to the fourth class that is termed *alternating* graph in the following. This scenario is illustrated by the black graph in Fig. 2 C. This characterizes a situation where a general statement about the conservativeness of statistical thresholds for likelihood ratios of the respective model parameter is not possible.

In fact, the appropriateness of the asymptotic approximation depends on the specific significance level for which e.g. a likelihood ratio threshold is derived. For this reason, the further analysis of empirical likelihood ratios λ from the parametric bootstrapping procedure covers both aspects: first, a classification of the *whole pp-plots graphs* in the presented four classes. And in a second analysis, an evaluation of the pp-plot graphs at certain *relevant quantiles*, i.e. for values of statistical thresholds for specific significance levels *α*. Such an example for 1 – *α* = 0.9 is depicted in Fig. 2 A, using the same classification areas as indicated in Fig. 2C for the whole pp-plots graph analysis.

### Benchmark models

For the following analysis, 19 nonlinear ODE benchmark models from the literature are used to investigate the empirical distribution of likelihood ratios. Table 1 summarizes all models, their properties and references.

**Table 1.** Table of all analyzed benchmark models.

| No. | Model name | $N_x$ | $r_x^{obs}$ (%) | $N_c$ | $N_{data}$ | $N_\theta$ | $N_{data}/N_\theta$ | Context | Reference |
| --- | --- | --- | --- | --- | --- | --- | --- | --- | --- |
| 1 | Alkan | 36 | 77.8 | 73 | 1733 | 44 | 39.4 | SB | [37] |
| 2 | Bachmann | 25 | 52.0 | 23 | 542 | 113 | 4.8 | SB<sup>†</sup> | [38] |
| 3 | Becker | 7 | 85.7 | 2 | 85 | 16 | 5.3 | SB<sup>†</sup> | [39] |
| 4 | Boehm | 8 | 62.5 | 1 | 48 | 9 | 5.3 | SB<sup>†</sup> | [40] |
| 5 | Brännmark | 9 | 66.7 | 8 | 43 | 22 | 2.0 | SB<sup>†</sup> | [41] |
| 6 | Bruno | 7 | 85.7 | 6 | 77 | 13 | 5.9 | SB<sup>†</sup> | [42] |
| 7 | Fiedler | 6 | 66.7 | 3 | 72 | 22 | 3.3 | SB<sup>†</sup> | [43] |
| 8 | Hass | 9 | 100.0 | 17 | 221 | 49 | 4.5 | SB<sup>†</sup> | [44] |
| 9 | Isensee | 25 | 64.0 | 109 | 713 | 46 | 15.5 | SB<sup>†</sup> | [45] |
| 10 | Lucarelli | 33 | 100.0 | 12 | 1755 | 84 | 20.9 | SB<sup>†</sup> | [46] |
| 11 | Merkle | 46 | 47.8 | 62 | 1141 | 197 | 5.8 | SB<sup>†</sup> | [47] |
| 12 | Kurzhunov | 3 | 33.3 | 1 | 46 | 9 | 5.1 | PK | [48] |
| 13 | Raia | 14 | 100.0 | 4 | 205 | 39 | 5.3 | SB<sup>†</sup> | [49] |
| 14 | Schwen | 11 | 54.5 | 7 | 292 | 30 | 9.7 | SB<sup>†</sup> | [50] |
| 15 | Sneyd | 6 | 33.3 | 8 | 135 | 15 | 9.0 | SB | [51] |
| 16 | Swameye | 9 | 33.3 | 1 | 46 | 16 | 2.9 | SB<sup>†</sup> | [52] |
| 17 | Tönsing - Flu | 3 | 33.3 | 1 | 14 | 5 | 2.8 | Inf | [10] |
| 18 | Tönsing - Zika | 8 | 12.5 | 1 | 58 | 9 | 6.4 | Inf | [10] |
| 19 | Zheng | 15 | 100.0 | 1 | 60 | 46 | 1.3 | SB<sup>†</sup> | [53] |

Models are numbered and abbreviated by the last name of the first author. $N_x$ : number of model states, $r_x^{obs}$ : fraction of observed model states, i.e. states that appear in the observation function, $N_c$ : number of experimental conditions, $N_{data}$ : number of data points, $N_\theta$ : number of estimated parameters, SB: Systems Biology model of cellular signaling pathways or gene regulation networks, Inf: Compartment model for infectious diseases, PK: Pharmacokinetic model, <sup>†</sup>: part of the benchmark model collection [54].

The collection comprises exclusively models that were published with experimental data and it covers a broad spectrum of modelled biological systems. Moreover, the models and their accompanied data sets differ widely in terms of size, e.g. number of estimated parameters or number of model states. Only the published data sets and the therein applied experimental design 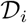, i.e. the choice of observables, of time points and of experimental conditions is used to simulate additional artificial, yet realistic data realisations as described earlier. All models share the characteristics of a partially observed nonlinear ODE system, i.e. they are calibrated using experimental data and thus are limited in the amount and quality of the finite data sample.

Some of the models exhibit a 100% fraction of observed states. These models should however also be considered as only partially observed, since for example not all states were observed also in all possible experimental conditions and perturbations of the biological system. Also, only a small subset of time points could be have been used for recording the data of certain model states. Despite the high fraction of observed states, the accompanied data sets of these models presumably do not contain the full information about the dynamical system. The ratio of the number of data points with respect to the number of estimated parameters lies between 1.3 and 39.4, i.e. at least as many data points are used as estimated parameters.

The selection of systems comprises 16 models which have a systems biology context, where mainly cellular signaling pathways are described and 14 of them originate from a comprehensive benchmark model collection [54]. In order to cover a broad spectrum of applications of dynamical models describing biological systems and to potentially reveal differences depending on the context, the set of systems biology related models is extended by two models that describe the dynamics of populations for infectious diseases taken from [10] and by a pharmacokinetic model of O_2_ concentrations in the human brain from [48].

Diverse model sizes with respect to their free parameters are considered. The smallest model has only 5 parameters to be estimated from 14 data points (17 Tönsing-Flu), while the largest model has approximately 200 fitted parameters estimated from 1141 data points (11 Merkle). A visualisation of absolute numbers of model parameters and the fraction of certain parameter types is given in Fig. 3 A. It appears that likewise to the broad range of model sizes, also the fraction of parameter types differs noticeably between the models.

**Fig 3.**
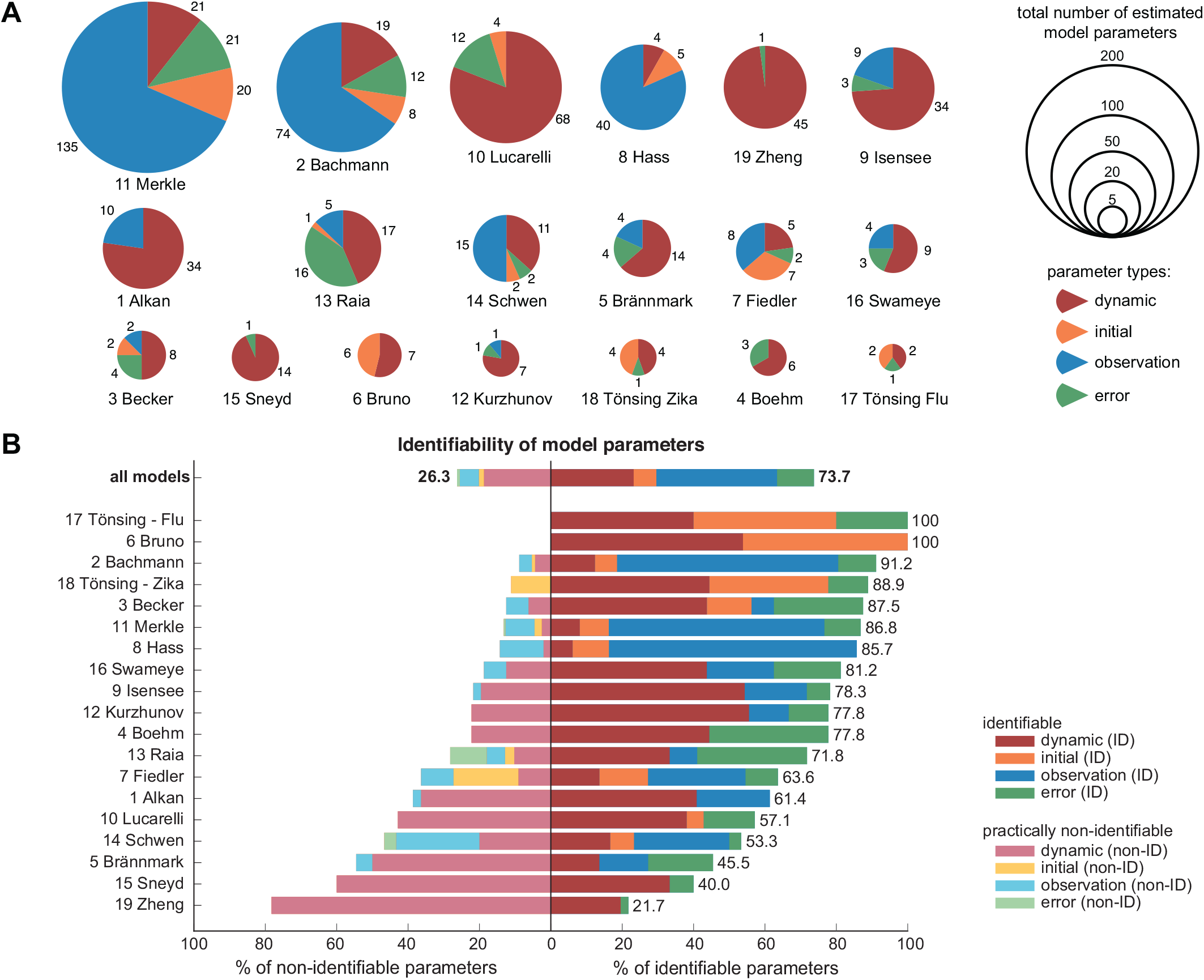
Overview of benchmark model properties and identifiability status. (A) Estimated parameters of all 19 models and distribution of parameter types. (B) Identifiability of model parameters based on the profile likelihood using the original data and a threshold of Δ_*α*=0.05_ = 3.84. Either identifiable parameters with finite profile likelihood-based confidence intervals (darker colors) or practically non-identifiable parameters (lighter colors) were identified. Structural non-identifiability was not observed.

The identifiability status of all benchmark model parameters is checked by calculating the profile likelihood for each model parameter, using the originally published data sets. For this analysis, the significance level *α* = 0.05, i.e. the commonly used threshold of Δ_*α*=0.05_ = 3.84 from asymptotic theory is utilized. Fig. 3B summarizes the results of the profile likelihood-based identifiability analysis.

Throughout all models, no parameter was identified as *structurally non-identifiable*, i.e. revealing a completely flat profile likelihood. A fraction of 26.3% of all parameters are however *practically non-identifiable*, all other parameters are *identifiable* with finite bounds of the profile likelihood. Likewise to the bare model properties from Table 1, a broad and evenly distributed spectrum appears in the fraction of practically identifiable parameters per model. Two models (17 Tönsing-Flu & 6 Bruno) are fully identifiable and most models have more than 50% identifiable parameters. Mostly dynamic parameters dominate the overall identifiability of the models, whereas initial value parameters, observation and error parameters are often non-problematic in terms of identifiability.

## Results and Discussion

### Observation: Empirical likelihood ratios

The empirical likelihood ratios of the parameters from the 19 benchmark models are analyzed in the following in order to check the validity of the asymptotic assumptions on their distribution. For all the in total 768 parameters, the bootstrapping procedure is employed and pp-plots of the resulting empirical likelihood ratios are analyzed according to the discussed classification scheme.

The presented procedure has large computational demands. First, the original best fit parameter set is used as initial guess for a single fit for each of the 19 models with each of the 500 data realisations. The main load however is generated at the second stage by the bootstrapping approach, which requires an additional loop of fits for each fixed parameter value and for each data realization. Especially large models with respect to the number of estimated parameters already require large amounts of computational resources for the single fits [54].

In total, the presented data sampling, bootstrapping and fitting procedure requires approximately 250 000 hours of single CPU time. All analyses were conducted using the Matlab-based *Data2Dynamics* modelling framework [55] on the *Baden-Württemberg High Performance Cluster (bwHPC) MLS&WISO*, where the analysis was completed within approximately 3 weeks of walltime by using up to 100 16-core 2.4GHz-nodes in parallel.

#### Analysis of whole pp-plots

In a first step, the *whole* pp-plots of all parameters are compared to asymptotic theory by assigning their graphs according to the discussed scheme to the four classes. The results of the classification for all parameters and separately for each parameter type are summarized in Fig. 4. Fig. 5 shows the pp-plots for all 784 parameters, grouped by the models and separated in identifiable parameters in the upper panel in Fig.5 A and practically non-identifiable parameters in the lower panel Fig.5 B.

**Fig 4.**
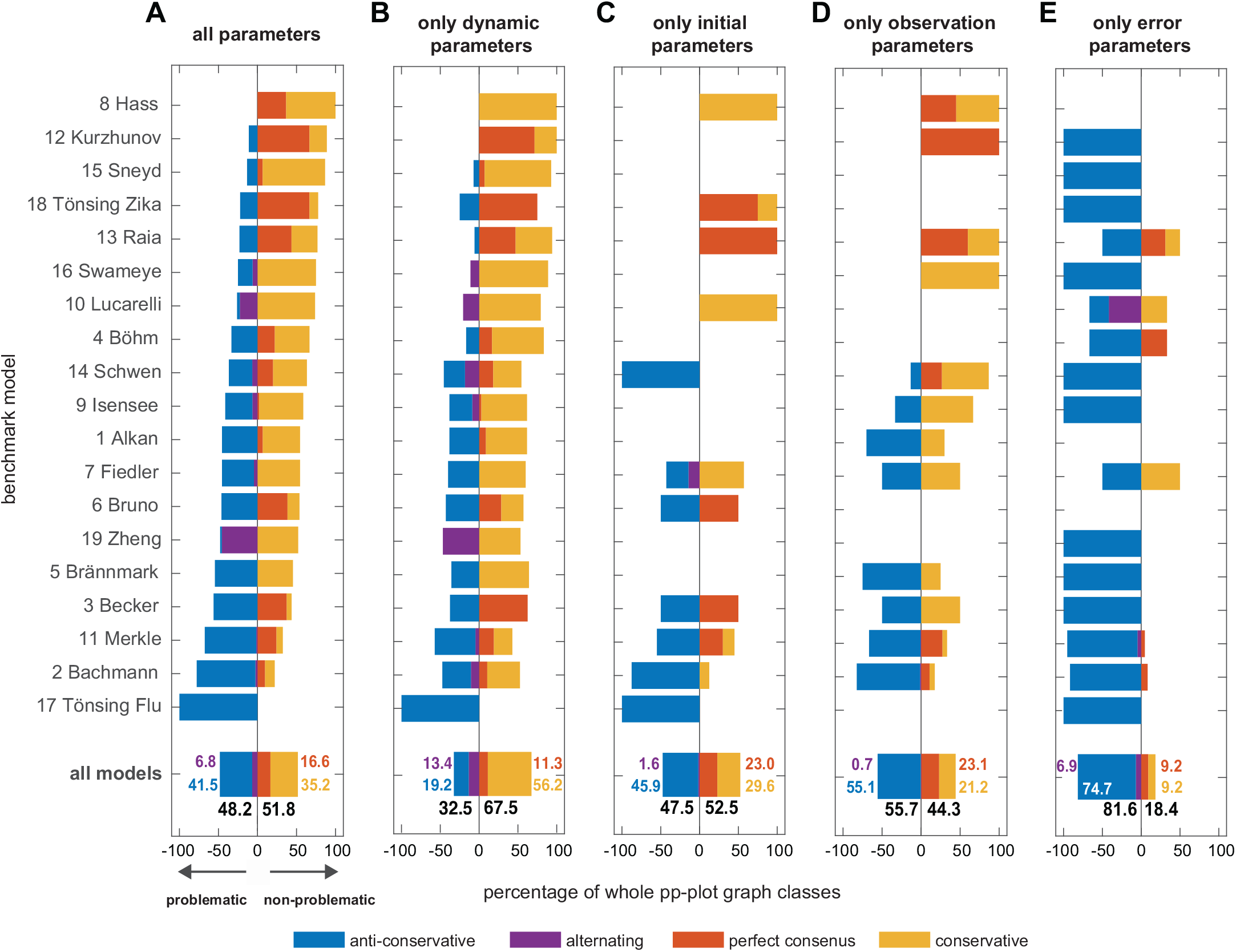
Classification results of whole pp-plots graphs compared to the 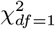-distribution. Results are sorted by the percentage of non-problematic, i.e. conservative and perfect consensus cases. The anti-conservative and alternating fraction of cases is indicated by a negative sign. (A) Overall results for all model parameters and (B-E) results separately for each model parameter type.

**Fig 5.**
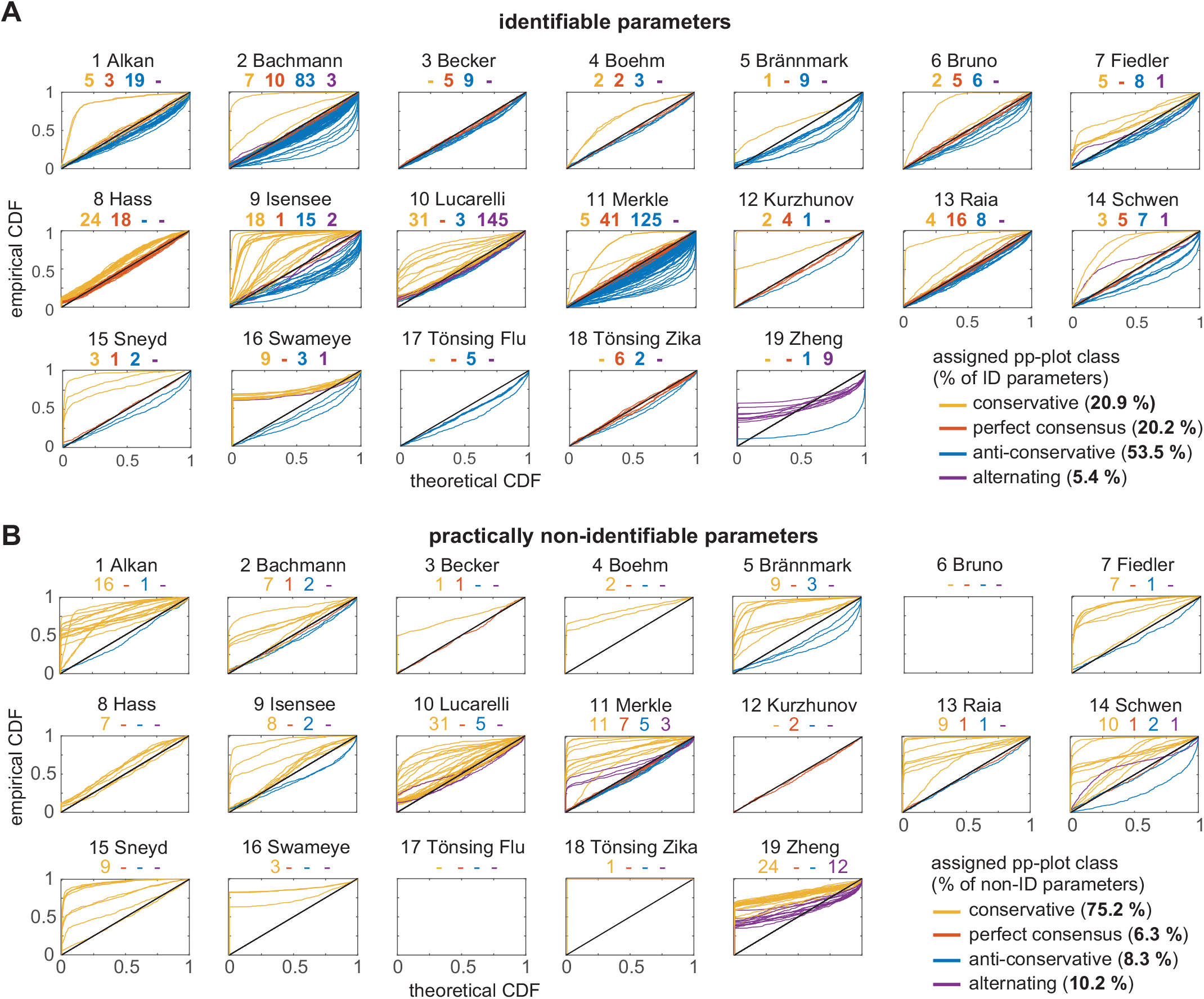
All pp-plots graphs of the estimated model parameters, separated by their identifiability status. (A) identifiable parameters and (B) practically non-identifiable based on their profile likelihoods. All graphs are classified in perfect (red), conservative (yellow), anti-conservative (blue) or alternating (purple) cases.

The analysis reveals a heterogeneous classification outcome throughout the different models, covering models with almost all parameters being in the conservative regime or showing a perfect consensus with theory(8 Hass), but also models with approximately 100% of anti-conservative cases(17 Tönsing-Flu). A surprisingly low overall fraction of only 16.6 % of the parameters in the perfect consensus class shows that only few empirical likelihood ratios are indeed 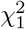-distributed. Only two models (12 Kurzhunov and 18 Tönsing - Zika) have more than 50% of parameters with a perfect overlap with asymptotic theory. The majority of analysed model parameters shows a distribution that is not in accordance with asymptotic theory. However, the *non-problematic* cases with *at least* conservative pp-plot graphs represent 51.8% of all analyzed parameters, shown as positive value bars in Fig. 4 A. Most models have a considerable fraction of anti-conservative or alternating pp-plot graphs. These *problematic cases* are grouped together with negative percentage in Fig. 4. In such cases it can not be guaranteed that the statistical thresholds for the likelihood ratio, e.g. for confidence intervals, are at least conservative. With 6.8% of all parameters, only a small fraction shows an alternating profile, i.e. with parts of the pp-plot graph in the conservative and in the anti-conservative part above or below the diagonal in the pp-plot.

An impression of the effect size of the pp-plot graph deviation from asymptotic theory is given in Fig. 6, which shows the average distance between the actual pp-plot graphs and the diagonal. For this, the vertical distance from to the graph to the diagonal is averaged over all 500 empirical likelihood ratios, i.e. data realisations for one parameter and normalized to the maximal possible deviation. The results are ordered with respect to the classification ranking for all parameters from Fig. 4 A. It shows a similar trend of the mean of the distances per model and the classification ranking, although the variance within the models is rather high. However, most pp-plot graphs show a relatively low distance to the diagonal, as indicated by the gray histogram on the right hand side in Fig. 6. If graphs show a large deviation from the diagonal, they tend to be more likely in the upper conservative region than in the lower anti-conservative region. In some cases in the conservative part, however, they reach the maximal value of the distance. Contrarily, in the more problematic case of pp-plot graphs in the anti-conservative region, they stay rather close to the diagonal in the perfect consensus region.

**Fig 6.**
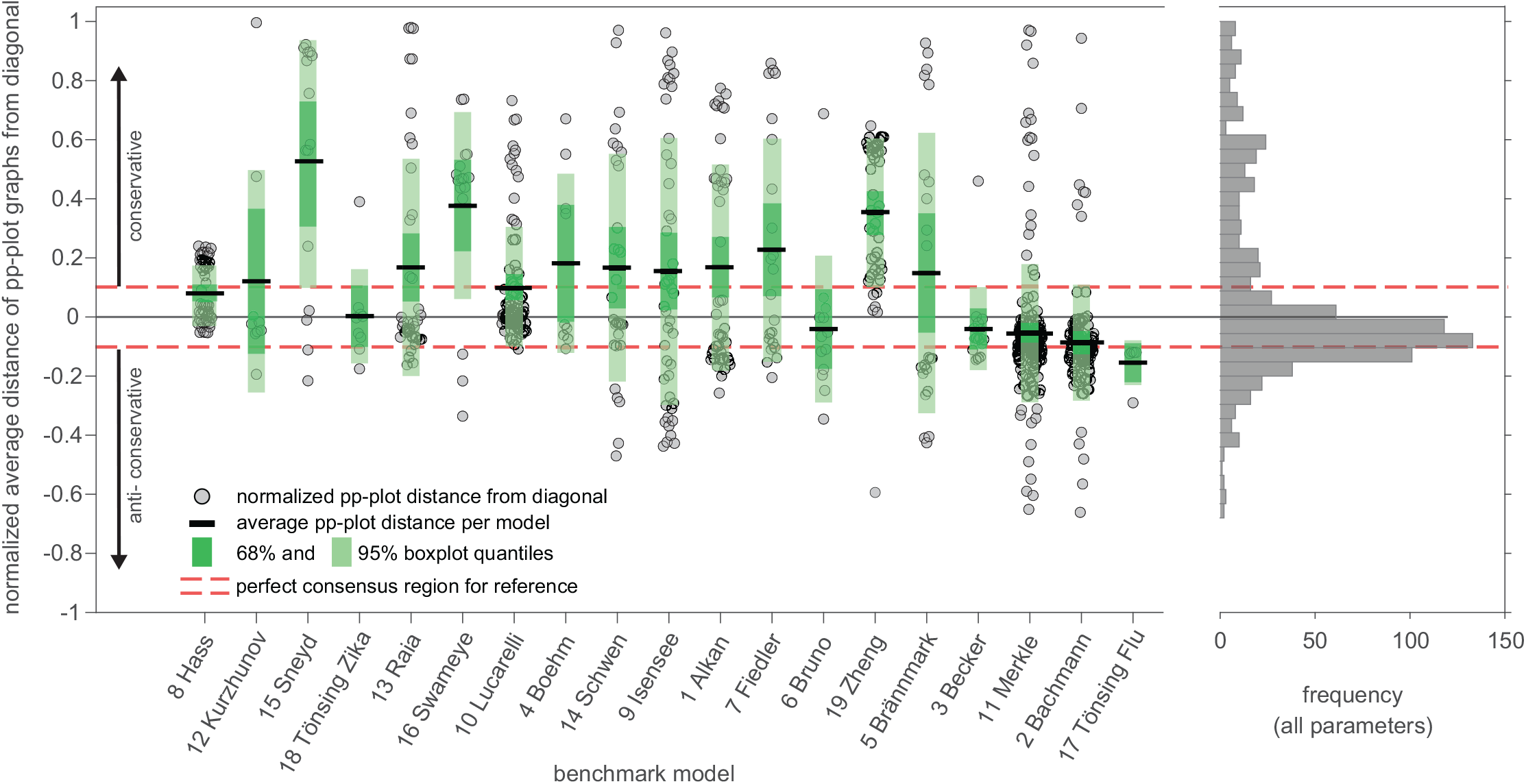
Average distance of pp-plot graphs from the diagonal. Distances are normalized by the maximal possible distance value, i.e. a graph lying on the upper x-axis at *p_emp_* = 1 for all *p_emp_* in the pp-plot. Models are ordered according to the ranking of the classification results from Fig. 4A. The histogram on the right shows the distribution of average distances from the diagonal summarized for all parameters.

When comparing the classification results for all parameters in Fig. 4A with the results for only dynamic parameters in Fig. 4 B, it appears that the partitioning of pp-plot graph classifications changes slightly, but would result in a similar ranking of models based on the fraction of non-problematic cases. Dynamic parameters dominate the overall classification of the models, but their non-problematic fraction is slightly higher. Initial value and observational parameters in Fig. 4C and D show a similar, yet less pronounced correlation with the overall results from Fig. 4 A. A comprehensive statement concerning the appropriateness of the asymptotic assumption for these parameter types remains vague, since many models do not have either initial value or observational parameters (cf. Fig. 3 A).

Error parameters are in most cases classified as anti-conservative, meaning that they fall in the problematic case where a statistical threshold for the likelihood-ratio test from asymptotic theory following Wilks’ theorem is too small compared to the empirical likelihood ratios from the bootstrapping procedure. It follows that, e.g. profile likelihood-based confidence intervals would be too small, regardless of the chosen significance level *α*. Thus, the uncertainty of the estimated error parameters would be underestimated and consequently the error model predicts the uncertainty to be smaller than it actually is. Interestingly, error parameters share the preferable property of being rather often identifiable which is desirable, but at the same time their pp-plots are mostly anti-conservative, which represents a rather problematic case.

A similar observation can be made concerning the relation between identifiability status and pp-plot classification, when observing at the actual pp-plots in Fig. 5 where identifiable and practically non-identifiable parameters are shown separately. Within the identifiable parameters in in the upper panel Fig. 5 A, all four classes are present, but only few models (2 Bachmann, 9 Isensee, 10 Lucarelli and 16 Swameye) show a considerable amount of more than five conservative cases (yellow lines). Contrarily, in the lower panel Fig. 5B showing the non-identifiable parameters almost all pp-plots indicate conservative pp-plot graphs. In contrast, anti-conservative parameters occur only in rare occasions for practically non-identifiable parameters (blue lines in Fig. 5 B), while almost every model exhibits at least one identifiable, yet anti-conservative parameter in Fig. 5 A.

Perfect consensus graphs are merely visible in Fig. 5 because these graphs are concentrated on a rather small region around the diagonal in the pp-plots. Absolute numbers are given for each class on top of the plots. Conservative and anti-conservative graphs are clearly visible, since they are spread over the whole plotting area. More precisely, it can be noted that the blue anti-conservative graphs touch the lower or right hand sided axes of the plots only in few cases. In contrast, the yellow graphs of conservative and non-identifiable parameters in Fig. 5B predominantly show a profile which instantaneously increases for p = 0 of the theoretical CDF indicating a rather large number of data realisations with very small empirical likelihood ratios close to zero.

#### Analysis of relevant quantiles

In practice, not all quantiles of the test statistic are relevant, since usually study results do not report the whole confidence distribution [56], but rather confidence intervals for a specific significance level *α* or confidence level 1 – *α*. Thus in this second analysis, the empirical likelihood ratios are examined only at specific thresholds values of the test statistic for typical assumed confidence levels. In contrast to the analysis of the whole pp-plot graph, here, only the specified confidence levels are employed in the pp-plot graphs, cf. Fig. 2 A. For this, the three classification regions from Fig. 2C are used, but only at the specific significance levels, corresponding to the respective quantiles of the test statistic. Consequently, the class of alternating graphs becomes obsolete, since at a specific level it is either in one of the non-problematic (perfect consensus or conservative) or in the anti-conservative region.

Fig. 7 shows the results for the frequently applied threshold of the test statistic for a significance level of *α* = 0.05. For this, the pp-plots of the empirical likelihood ratios are classified only at the confidence level of 1 – *α* = 0.95. Specifically, it is assessed whether the calculated statistical threshold 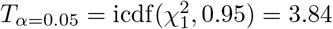 of the test statistic’s distribution under the asymptotic assumptions holds by checking if at least 95% of the empirical likelihood ratios from the bootstrapping procedure are smaller than 3.84. This is done by checking, if the pp-plot graph of the empirical CDF at *p_theo_* = 0.95 lies within the perfect consensus region, below or above it. Compared to the results from the *whole* pp-plot analysis (cf. Fig. 4), the analysis at the 0.95-quantile in Fig. 7 shows a similar ranking of the models, although considerably more parameters reveal the preferable perfect consensus case. However, the overall amount of anti-conservative cases remains at a similar level and most models again show a heterogeneous pattern, with in the majority of models at least one third of as anti-conservative classified parameters.

**Fig 7.**
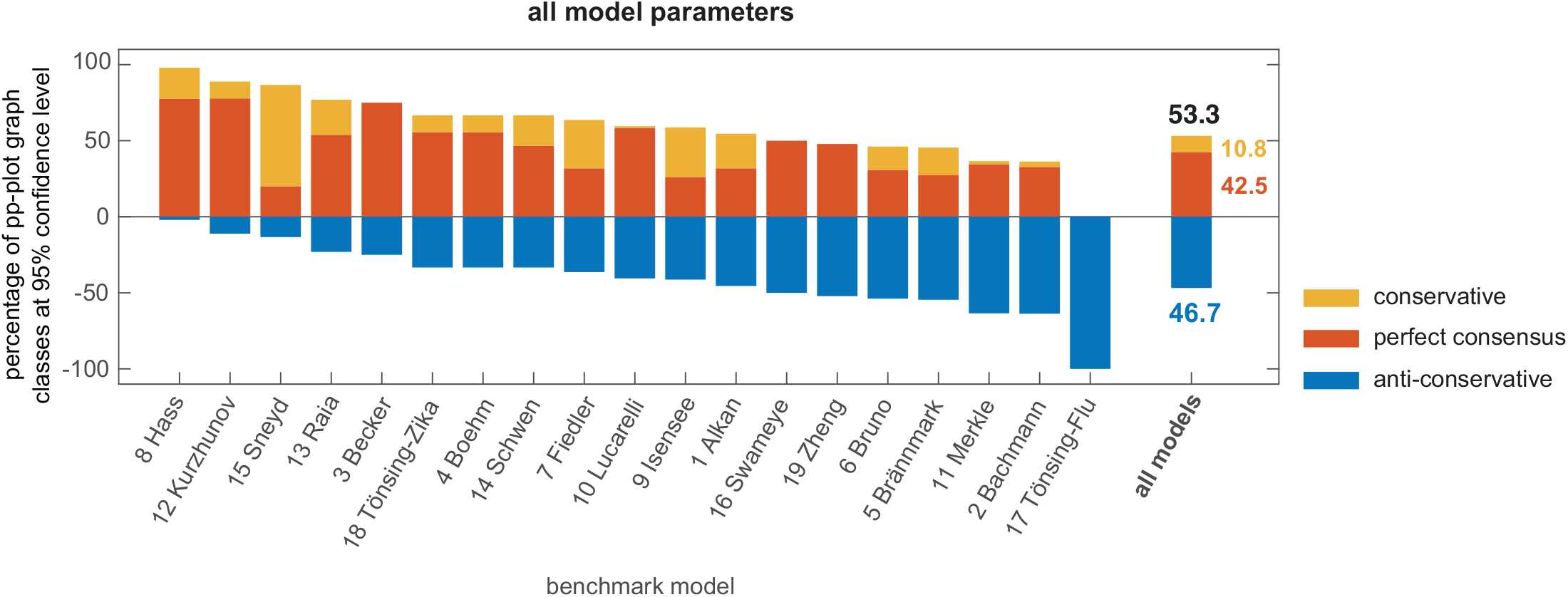
Classification results of pp-plots graphs at the 95% confidence level. Results are sorted by the percentage of non-problematic, i.e. conservative and perfect consensus cases. The model ranking remains similar to the whole pp-plot graph analysis (cf. Fig. 4)

The same kind of analysis is repeated for a set of specific confidence levels, in order to check if the appropriateness of the asymptotic assumption improves when considering the whole range of possible confidence levels, including the prominent 68% and 99% levels. Since the ideal situation of mostly perfect consensus cases is not reachable anyhow, the following analyses examine non-problematic cases, i.e. perfect consensus *and* conservative cases versus the problematic anti-conservative parameters. For this, the conservativeness ratio

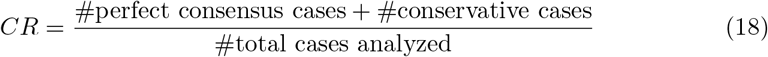

is defined, which serves as a measure for the appropriateness of the assumed test statistic distribution.

Fig. 8A shows the conservativeness ratio *CR* for a range of confidence levels for all analyzed parameters of the benchmark models, separated in their parameter type, their identifiability status and for the individual models. A *CR*-value close to 1 indicates a good asymptotic approximation of the likelihood ratio statistic at the respective confidence level. In the heatmap in Fig. 8A it can be observed that the *CR* indeed varies with the confidence level of the test statistic’s threshold. As a general trend, it can be concluded that lower confidence levels show higher values of the *CR*, meaning that they are more often in accordance with asymptotic theory than higher confidence levels.

**Fig 8.**
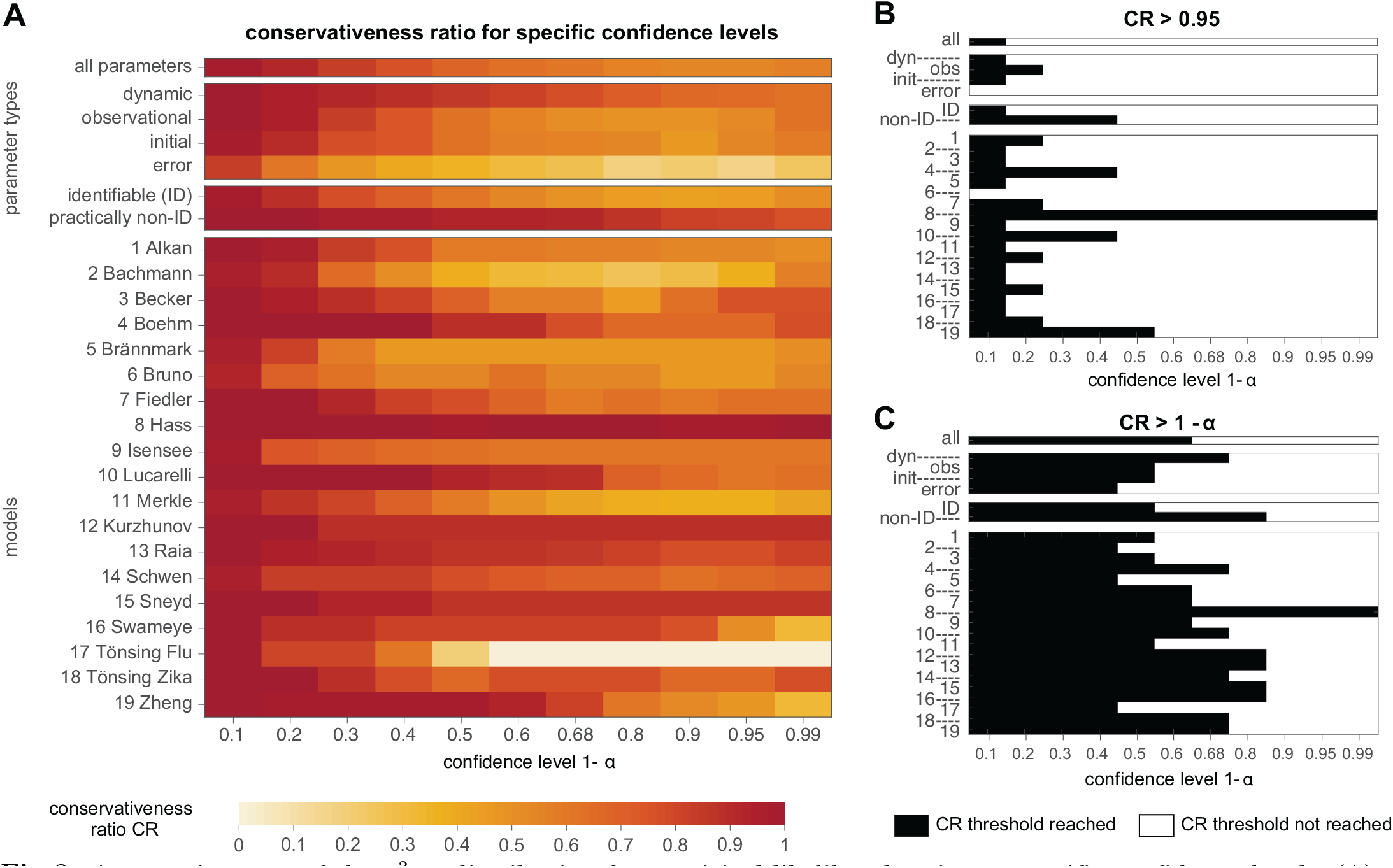
Appropriateness of the 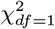-distribution for empirical likelihood ratios at specific confidence levels. (A) Heatmap of conservativeness ratio *CR*, i.e. fraction of non-problematic cases in the respective parameter group or model. (B) Parameter groups or models with *CR* larger than 95% at the confidence level are indicated by black tiles in the upper panel. (C) Less strict requirement assessed by an *CR* of at least 1 – *α*% at the respective confidence level.

For a binary decision whether the approximation of the empirical likelihood ratios by the assumed 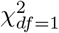-distribution holds or if it is violated, two thresholds for the CR are defined. One typical requirement would be that the asymptotic approximation should be valid in at least 95% of the cases, regardless of the considered confidence level. The heatmap in Fig. 8B indicates that this requirement of *CR* ≥ 95% is fulfilled for almost no model at any confidence level. Precisely, only for a very small confidence level of 10%, at least 95% of the model parameters show a sufficient accordance with asymptotic theory. Of course, the formulation of such low confidence levels is troublesome, as it corresponds to a not desirable type I error of e.g. 90 %. Two exceptions are model ‘6 Bruno’, which does not reach a *CR* ≥ 0.95 for any of the analyzed confidence levels and model ‘8 Hass’, which exhibits the opposite case of a sufficient *CR* ≥ 0.95 for all analyzed confidence levels.

Another, yet weaker requirement for a sufficient *CR* is based on the same type I error at all stages of the analysis, i.e. for the likelihood ratio statistic, the respective confidence intervals, and for the appropriateness of the choice of the 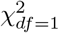-distribution. With this definition, a *CR* ≥ 1 – *α* would be a sufficient condition for a valid asymptotic assumption, since for a e.g. 68 %-confidence interval, only 68% of the empirical likelihood ratios need to be non-problematic. Fig. 8C shows the results for this criterion applied on the heatmap from panel A. It appears that considerable more confidence levels are in accordance with this criterion than with the stricter rule shown in panel B. However, already for a confidence level 1 – *α* ≥ 0.5, many groups or models do not have a sufficiently large CR and thus, the validity of asymptotic approximation from Wilks’ theorem cannot be guaranteed.

These results confirm what was already concluded from the previous analysis of the pp-plots: The validity of the asymptotic assumption in Wilks’ theorem applied to realistic ODE models cannot be guaranteed, neither for the whole pp-plot, nor for typically utilized quantiles of the distribution, such as e.g. 0.68, 0.9, 0.95 or even 0.99. By comparing the *CR* results for the error model parameters with the average *CR* or other parameter types, it appears that they show considerably less agreement with the *χ*^2^-distribution. More precisely, error model parameters rather exhibit the anti-conservative case, in which confidence intervals tend to be too small. Furthermore, also the group of identifiable parameters are more often problematic, when compared to the CR of practically non-identifiable parameters. Interestingly, practically non-identifiable parameters rather exhibit the conservative case, i.e. confidence intervals tend to be too large (see also Fig. 5). Looking at the individual models, the analysis reveals a large heterogeneity in the *CR*-values. There are models that show a good agreement of the asymptotic assumption (e.g. model 8 Hass) over all quantiles, or at least for quantiles up to 0.68 or 0.8. But it can be observed that there are also models where the asymptotic assumptions even for a confidence level of 1 – *α* = 0.5 holds at least conservatively in only 50% of the cases. It should be further noted as conclusion of the analysis at the relevant quantiles, that approximately 50% of the parameters are classified as anti-conservative at the commonly utilized confidence level of 0.95, cf. Fig. 7.

### Cause: Data space interpretation

To motivate a possible explanation, how the different classification results and pattern in the pp-plot graphs are related to the model and data properties, a geometrical interpretation of parameter estimation utilizing the concept of the data space is introduced in a short digression. For linear models, fitting a model to the data is typically realized by using the pseudo-inverse of the design matrix [57]. The pseudo-inverse can be interpreted as a projection operator which minimizes the Euclidean distance between the so-called model manifold and a point in the so-called *data space*.

The data space for *n* data points is represented by the *n*-dimensional space 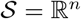, where each data point corresponds to one dimension of 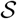. Thus, one data set, i.e. one specific noise realization of all data points corresponds to a single point in the n-dimensional data space. Multiple data realisations of the same experimental design 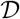 can be displayed side by side in the data space by individual data space vectors. Contrarily, an additional measurement which extends the data set, i.e. for an alternative experimental design 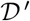 would introduce an additional dimension to the data space 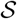.

The model in the data space is represented by a manifold 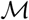 with dimension *m* ≤ *n*. Residuals *res_i_*(*θ*) = *y^data^*(*t_i_*) – *y^model^*(*t_i_, θ*) quantify the deviation of the observation *y^data^* from the model prediction *y^model^* at time point *t_i_* for a parameter vector *θ*. The objective function value

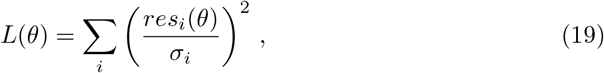

i.e. the sum of squared weighted residuals, corresponds to the Euclidean distance in the data space between data point and model manifold. The standard deviation *σ_i_* thus corresponds to a re-scaling of the respective data space axis. For an additive Gaussian error model, the re-sampled data is distributed spherically in the *n*-dimensional data space. Thus, the data realisations are centered around the true parameter value on the model manifold. Model fitting to such a data realization is the local orthogonal projection on the closest tangent of the model manifold. In other words, parameter estimation is equivalent to finding the minimal Euclidean distance between data and model in the data space. Differences in terms of the objective function are reassembled by the Pythagorean theorem [58, 59].

Fig. 9A shows an illustrative example of a one-dimensional model with one model parameter *θ* and with two observations *t*_1_ and *t*_2_. The model predictions at *t*_1_ and *t*_2_ can be interpreted in the two-dimensional data space by

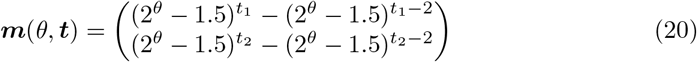

yielding a nodal curve as model manifold (motivated by [60]). Interpreted as function of the true parameter value *θ*, the model manifold yields different local shapes, which have characteristic consequences for the model fitting, as the shape of the manifold influences the distances between data and model prediction as discussed in the following. For the illustration, four scenarios are defined by four different values of the model parameter *θ*, indicated by a blue cross on the model manifold in Fig. 9 A,D,G,J. Data is generated by sampling from an additive Gaussian error model, yielding data realisations in the data space that are indicated by gray crosses. A fit of the model to such a data point is an orthogonal projection of the data point in the data space on the model manifold, shown as red circles. Analogously to the likelihood ratio or the differences in terms of the log-likelihood (cf. Eq. (8)), distances of the model fits (red circle) to the location of the model prediction for the true parameter value, i.e. maximum likelihood estimate (blue cross) are displayed in the histograms in Fig. 9 B,E,H,K. If the asymptotic assumptions about the likelihood ratios hold, these distances are expected to be 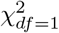-distributed following Wilks’ theorem. As a reference, a 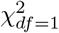-like distribution is depicted in the histograms as a red line. The distances in the data space, i.e. the analog to empirical likelihood ratios for different data realisations are compared against the expected 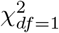-distribution in pp-plots in Fig. 9 C,F,I,L according to the previous analysis and classification of empirical likelihood ratios from the bootstrapping procedure for nonlinear ODE models.

**Fig 9.**
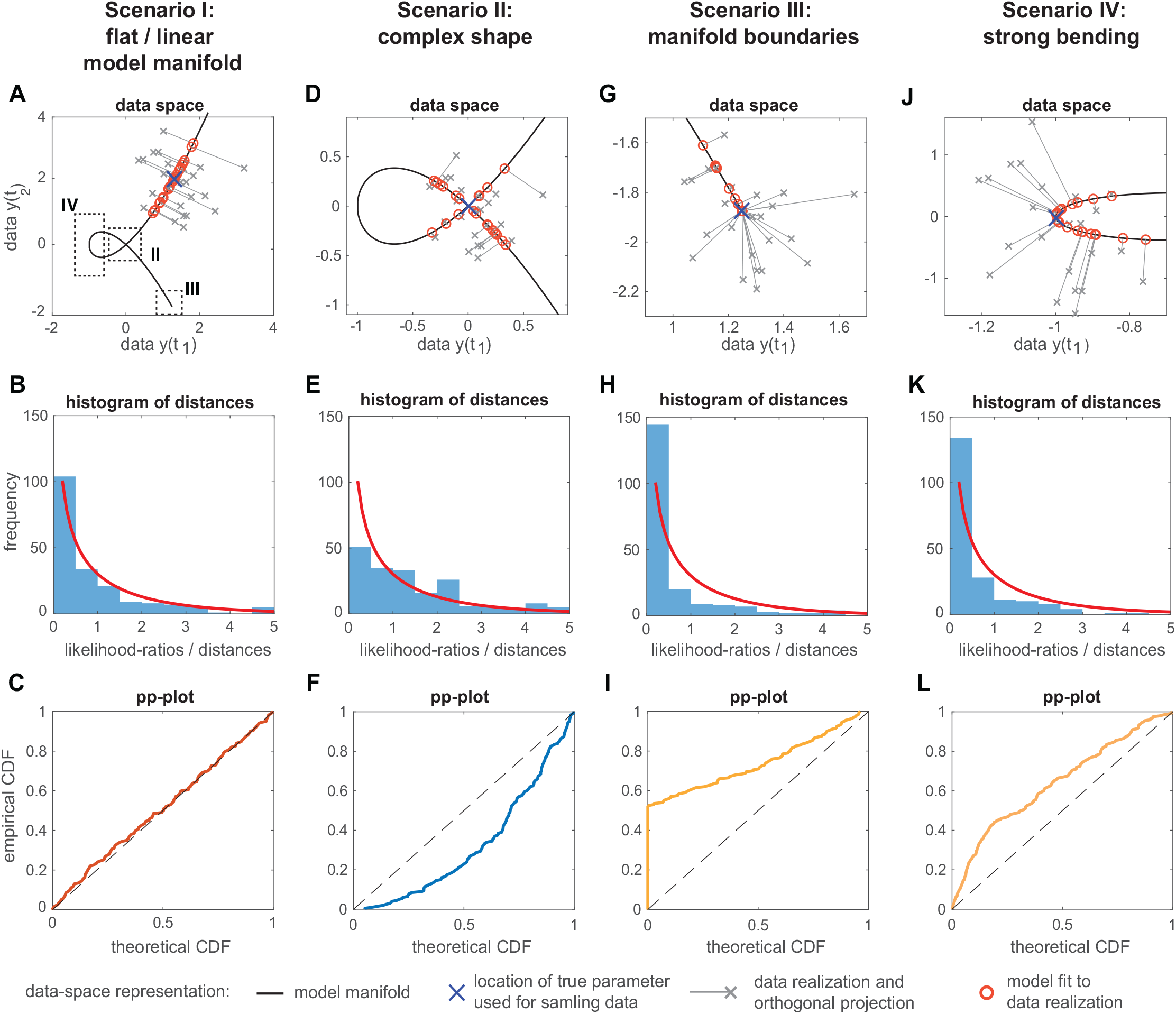
Data-space representation of four different scenarios using the nodal curve model manifold example. (A,D,G,J) Different shapes of the model manifold (black solid line) in the data space with a choice of data realisations (*N* = 25 out of 200, gray crosses) drawn from a n-ball around the true parameter (blue cross) and fit onto the model manifold (gray solid line and red circles). Distances between the true parameter and the fits (*N* = 200) correspond to the empirical likelihood ratio. (B,E,H,K) Corresponding histograms to each scenario illustrate the distribution of the distances, i.e. likelihood ratios compared to a 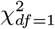-distribution (red line). (C,F,I,L) The pp-plot panel allows for a detailed comparison of the empirical likelihood ratios to a 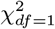-distribution (*N* = 200).

In the first scenario I, the local shape of the model manifold, i.e. the surface region populated by the single fits (red circles), is almost uncurved and thus, the outcome is similar to a linear model. In higher dimensions such a model manifold would be a flat *n* – 1-dimensional hyperplane. In accordance to asymptotic theory, the empirical objective function distances are indeed 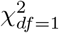-distributed, as verified by a *perfect consensus* graph in the pp-plot, almost identical to the diagonal in Fig. 9 C.

Further increasing the value of the true model parameter *θ* moves the point from which the *N* = 200 data realisations are sampled towards the intersection of the nodal curve (scenario II), cf. Fig. 9 D. As a consequence, the estimated model parameters populate a more complex surface of the nodal curve when being projected to the model manifold in vicinity of the intersection. Since in this situation the model manifold lies more densely in the data space, the probability of having the manifold in close vicinity around an individual data realization is higher than for the linear model case. Compared to the flat manifold from the first scenario however, the distribution of distances between individual fits and the true parameter on the model manifold changes towards more increased distances, while the distances between data realisations and individual fits tend to decrease. The effect is depicted in the histogram in Fig. 9E that differs from the exponential decay-like shape in scenario I, as it is more uniformly distributed. As a consequence, the pp-plot unambiguously reveals an anti-conservative graph, i.e. indicating that a statistical threshold for a test statistic distributed as 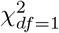 would be too small compared to the actual empirical distribution, which contains more larger distances.

Scenario III illustrates the effects of boundaries at the model manifold in the data space, which can be caused by two conditions. Either it originates from boundaries of predefined parameter constraints or from boundaries caused by the model’s nonlinearity. The latter case appears for example when increasing or decreasing the value of a model parameter or a combination thereof does not change the model output at a specific data point. Such a scenario might appear even in simple models. Thinking of an exponential decay *f* (*κ, t*) = *e^-κt^* or a saturation dynamics *f* (*κ, A*_0_, *t*) = *A*_0_(1 – *e^-κt^*), where for example the value of the functions at a certain time point is bounded, but a noisy data point might lay beyond this boundary, of the model manifold. For instance, the model prediction for the exponential decay is bounded to values larger than zero, whereas the trajectory of the saturation dynamics does not exceed the saturation level *A*_0_, even when tuning the parameter *κ* to extreme high values. Such effects can cause a practical non-identifiability, where the parameter *κ* might only be constrained in one direction by the data, i.e. its profile likelihood exceeds a Δ_*α*_-threshold only for one tail of the profile. However, tuning the parameter to more extreme values on the other side of the profile does not change the discrepancy between data and model prediction, i.e. the objective function and thus the profile likelihood stays constant without hitting the Δ_*α*_-threshold.

The part of the model manifold selected for Fig. 9G shows this setting of a parameter at the boundary of the manifold. As the value of the model parameter *θ* from Eq. (20) is increased, the model trajectories of the two considered data points *y*(*t*_1_) and *y*(*t*_2_) will not exceed the point ***m**^bound^* = (1.25, −1.875) in the data space, although noise realisations of the data may lie behind this point. Two groups of data realisations can be identified for scenario III in Fig. 9 G. Data points of the first group in the upper left part are projected on the linear-like part of the model manifold, similar to the case of the almost uncurved model manifold in Scenario I. These fits result in estimated parameters values (red circles) with individual and finite distances to the true parameter at the boundary (blue cross). On the contrary, all data points from the second group in the lower right region of the data space are projected onto the same point of the model manifold, which coincides with the model manifold’s boundary *_m_^bound^*. In other words, all these data points imply a model fit with the same model prediction (blue cross), but different objective function values (gray lines). In addition, all these model fits lie on the same point in the data space, but do not restrict the model parameter *θ* to upper values, since all parameter values beyond this value yield the same model prediction, as already discussed e.g. for the saturation dynamics with a very large parameter *κ*. This characterizes a practical non-identifiability. All fits of this second group share a zero value for the distances between the fitted parameter value and the true parameter value. The mismatch to the assumed 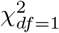-like distribution of these distances is reflected by a huge peak in the bin distances at zero in the histogram in Fig. 9 H. Correspondingly, the graph of the pp-plot in Fig. 9 I shows a sudden increase for a probability *p* ≈ 0 of the theoretically expected CDF, similar to the typical shape of the observed non-identifiable conservative parameters in Fig. 5B in the previous analysis. Here, a statistical threshold based on asymptotic theory would be too large in comparison to the empirical bootstrapping sample, i.e. it would be classified as conservative.

Strong bending of the model manifold is shown in Fig. 9 J for Scenario IV, where the true model parameter is located in the turning point of the nodal curve at the most left-handed side of the data space. To illustrate the strong bending effect using the model from Eq. (20), the *t*_2_ axis is squeezed by using a smaller *σ*_2_ for the additive noise. Likewise to scenario III, roughly two groups of data points can be identified, while their separation line is rather a transition region and less pronounced than in the previous scenario. Data points with *y*(*t*_1_) > −1 have a projection direction almost parallel to the *t*_2_-axis and roughly reassemble the fit to a flat model manifold as in scenario I. The remaining data points with *y*(*t*_1_) < −1 are projected to roughly the same region on the model manifold in very close vicinity of the turning point of the model manifold, where the true model parameter is located. Thus, distances between the model prediction of the estimated parameters for the latter group of data points and the model predictions of the true parameter value are smaller than expected from asymptotic theory. This discrepancy is similar to scenario III, but less extreme, as can be seen from the histogram and 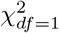-reference in Fig. 9 K. In the same fashion, the pp-plot graph reveals a less extreme conservativeness, i.e. it clearly deviates from the diagonal, but does not show the sudden increase for lower probabilities of the theoretical CDF. This indicates, that the bending is not as extreme, so that the data points are not projected exactly on the same model manifold turning point, but small distances to the true parameter value on the model manifold occur. This effect would change for the case of a more extreme curvature, where the upper and lower arm of the manifold in Fig. 9 J would almost fall together. The resulting model manifold would look like a single flat line with a boundary similar to the manifold from scenario III. In such a case, all left-handed data points would be projected to exactly the same point, yielding a pp-plot graph similar to the Fig. 9 I. It should be noted, however, that this case does not represent a model manifold boundary for practically non-identifiable parameters, whose value can be increased without changing the objective function. In fact, such a case would yield a single point estimate and a profile likelihood of the estimated parameter with a unique minimum and finite profile likelihood-based confidence intervals, which would be conservative, though. These extreme curvatures, which might be even more pronounced for models with more parameter dimensions and in a high-dimensional data space, help to explain why identifiable parameters show pp-plot graphs similar to Fig. 9 L, although they are not located at a model manifold boundary as in scenario III.

### Cure: Adapted statistical thresholds

#### Impact of increased data sample size

After observing the deviation from asymptotic theory in a considerable fraction of cases and having formulated a geometrical explanation of its origin, three approaches handling the finite-sample case are discussed. First of all, the transition from the finite-sample case towards the asymptotic limit is inspected, showing on the one hand that indeed the amount of information contained in the data ist the reason of the deviation and on the other hand highlighting that increasing the number of data points in realistic applications does cure the issue.

To this end, the impact on the pp-plots of an altered experimental design 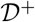 with additional data points in the benchmark models is investigated The original experimental design 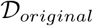 is extended by simulating additional data data points for time courses of observables which were already contained in the original design, for example by doubling the temporal sampling. Fig. 10 shows the results of two experimental designs 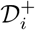 with increased temporal sampling for three models and compares the results to the original design.

**Fig 10.**
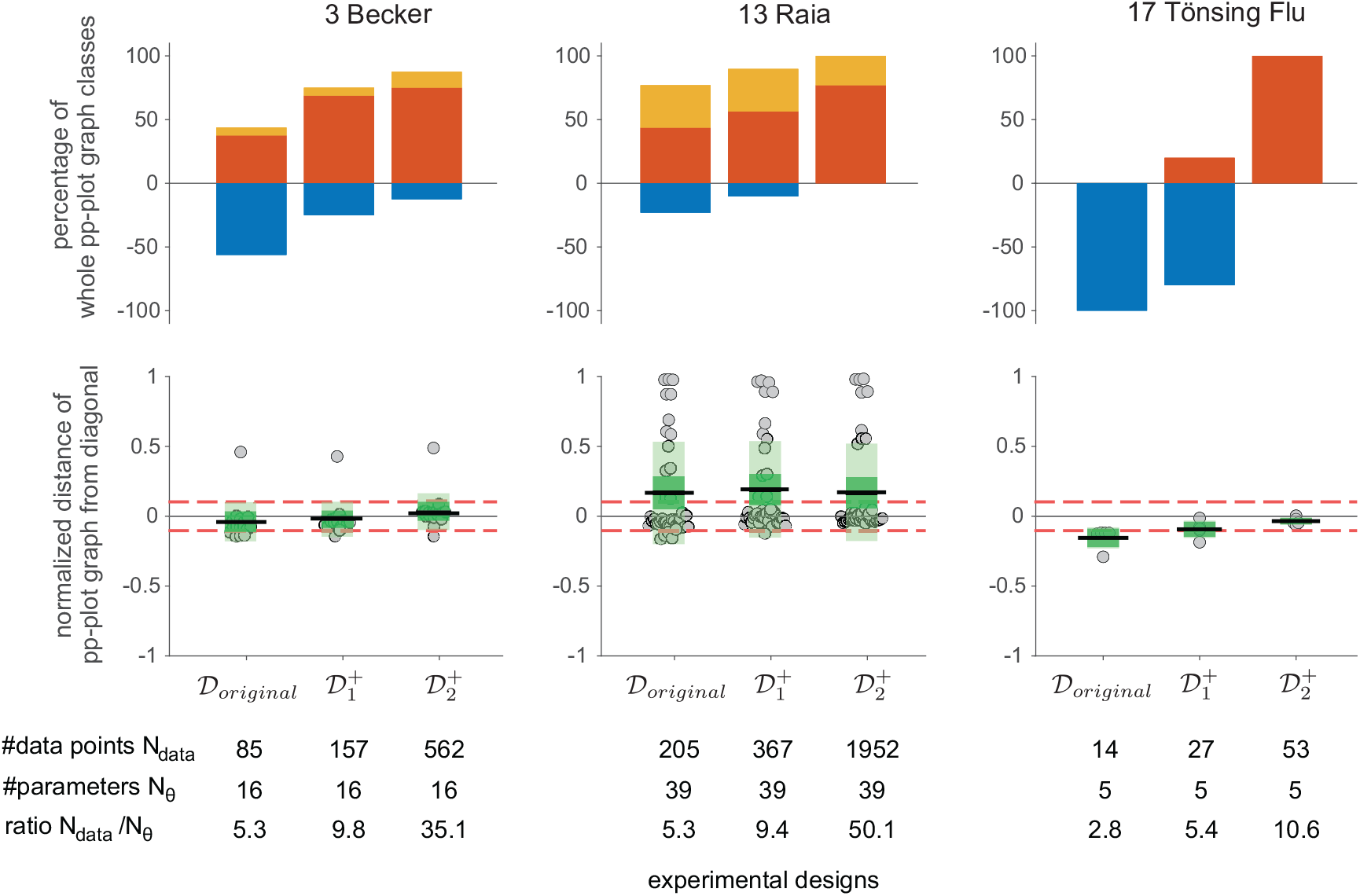
Impact of experimental designs with increased amounts of data. Whole pp-plot graph classification results and average normalized distances from the pp-plot graphs to the diagonal for three illustrative benchmark models. The outcome for the originally published design 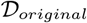, and two artificial designs 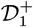 and 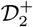 with increased temporal sampling is shown. The respective number of estimated parameters, total number of data points and their ratio is given below.

By shifting the design towards the asymptotic case, i.e. by increasing the number of data points, it can be observed that the classification of the whole pp-plot indeed tends to a higher number of parameters with non-problematic cases for all three models. For the model ‘17 Tönsing-Flu’, the classification even changes from completely anti-conservative to desirable 100% perfect consensus. It can be further observed for the models ‘3 Becker’ and ‘13 Raia’ that the average distances from the diagonal change only slightly, while the classification result indicates a clear improvement of the appropriateness of the asymptotic assumptions. However, a certain amount of conservative and anti-conservative cases persists, even for large amounts of data, compared to the original design. In these cases, the asymptotic regime is not yet reached by increasing the temporal sampling of the existing observables, as performed here. Extreme amounts of data points or the incorporation of more data points from additional observables, that carry more relevant information about the systems, would yield a more suitable *asymptotic experimental design*, cf. [61, 62]. However, in typical systems biology related projects simply increasing the sample size is not a viable option since data generation in molecular biology is often laborious and costly. Changing the experimental design by including additional observables might also be impossible for example due to missing antibodies for certain compounds. Therefore, we discuss in the following two different approaches to cope with the finite-sample case when analyzing likelihood ratios.

#### Bartlett correction by extensive search

In order to correct for the deviation of the likelihood ratio distribution from the asymptotic limit, the Bartlett correction can be used. Although it is not possible to calculate the appropriate correction factor in advance from the model properties in realistic applications of nonlinear ODE models investigated here, an approximate can be derived based on the results from the bootstrapping procedure. For this, the previous analysis of pp-plot classes at the 95 %-quantile is repeated but with altered empirical likelihood ratios

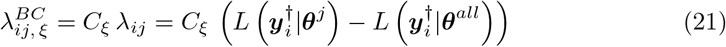

using a set of 600 Bartlett-like correction factors *C_ξ_* ∈ [0.1,…, 1,…, 600], cf. Eq. (13) and (16). Corresponding pp-plots are calculated, evaluated at the 95%-quantile and assigned to either the perfect consensus, conservative or anti-conservative case, according to the previously discussed scheme. Despite the large number of results from the 600 correction factors for each of the 784 parameters, this enables for a relatively cheap exhaustive search for an optimal Bartlett-like correction factor *C_ξ_*.

For all except one parameter, at least one individual Bartlett-like correction factor *C* can be found within the set of *C_ξ_* ∈ [0.1,…, 1,…600], yielding a *perfect consensus* pp-plot at the 95 %-quantile. Only in one special case, a parameter from the model ‘18 Tönsing-Zika’, which implements prior knowledge information, is assigned to the conservative case for all applied corrections factors even in an extended range of 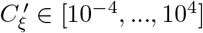. As expected from the previous analysis (cf. Fig. 7), the procedure yields approximately 41.2% of the parameters revealing a perfect consensus for uncorrected likelihood ratios with factor *C* = 1. For the remaining parameters, the individual *optimal* correction factor is defined as the *C_ξ_* ≠ 1 closest to 1, that yields a perfect consensus pp-plot at 95 %.

Fig. 11 A shows the 95 %-pp-plot classification result as coloured tile for all correction factors *C_ξ_* as indicated on the y-axis and for all analyzed parameters shown on the x-axis. These parameters are sorted by their individual optimal correction factor as indicated by the black line in the heatmap. Thus, each column shows the effect on the pp-plot classification for the specific model parameter with varying correction factors. It appears that the Bartlett correction affects the classification results in a non-monotonic way: the non-problematic perfect consensus and conservative cases fluctuate for smaller correction factors, whereas there is a sharp separation between the problematic cases and the anti-conservative classification. When sorting the parameters with respect to their optimal correction factors, an approximative conservativeness ratio (*aCR*) for a specific correction factor appears from the percentage of parameters on the right hand side of the intersection between the specified correction factor and the black line of optimal correction factors. For example, the white horizontal line in Fig. 11A indicates the uncorrected scenario with correction factor *C* = 1. It intersects with the black line of the parameters’ optimal correction factors and a conservativeness ratio *CR* of 52% can be read off the x-axis, since this fraction of the parameters reveal a perfect consensus or a conservative classification, whereas most of the parameters at the right hand side of the intersection are classified as anti-conservative. Exceptions from the anti-conservative classification on the right hand side occur when the classification is non-monotonic with respect to the correction factor, i.e. when a parameter is classified as *conservative* for the respective correction factor (*C* = 1), but the optimal correction factor for a *perfect consensus* is larger. For this reason, only an *approximative* conservativeness ratio aCR is available in this visualisation from the projection of the intersection on the x-axis of the heatmap. It slightly differs from the previously calculated conservativeness ratio CR as for example in Fig. 7.

**Fig 11.**
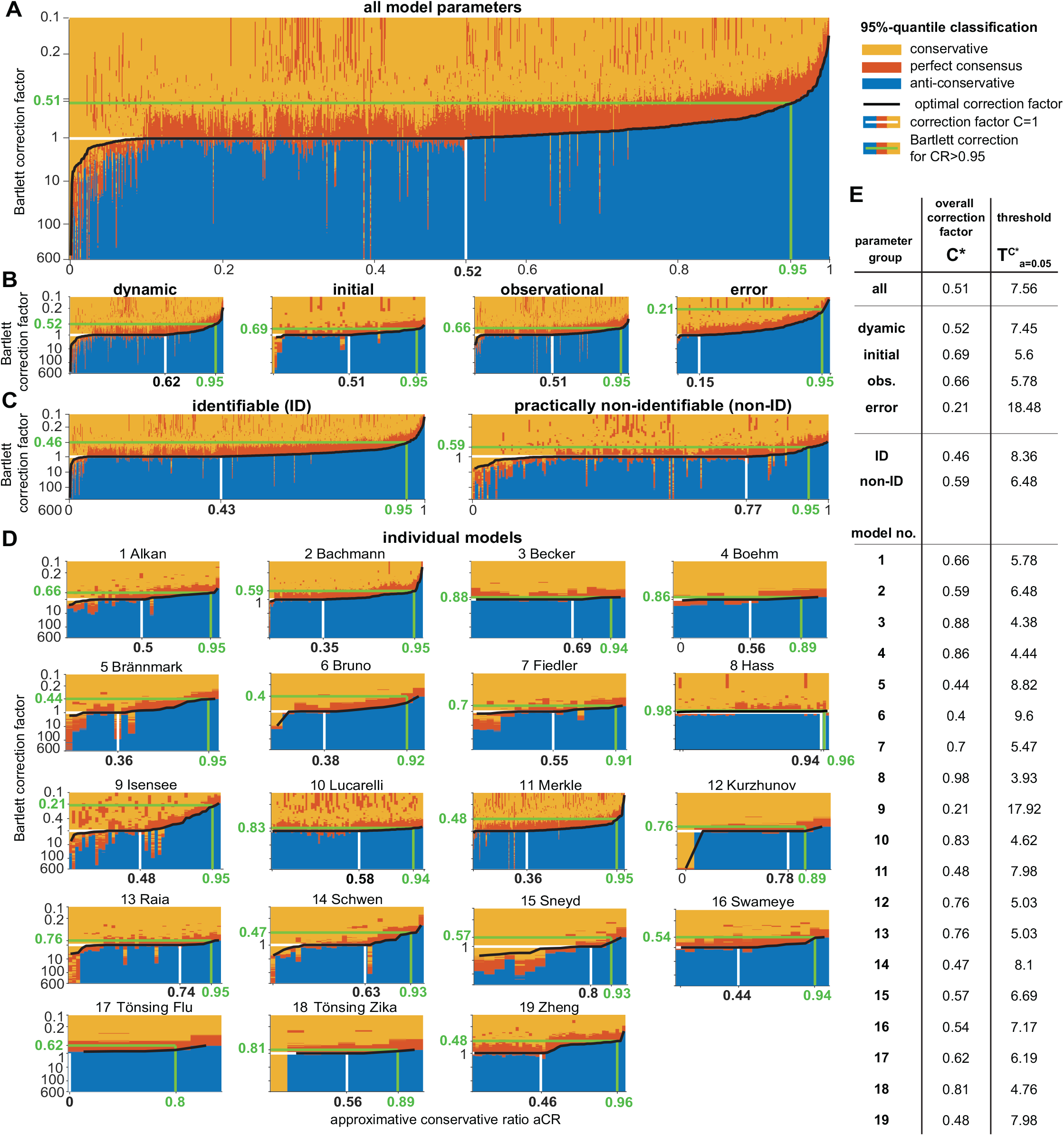
Bartlett correction results of pp-plot classifications at the 95% confidence level. Heatmap colors according to pp-plot classification for each Bartlett correction factor as indicated on the y-axis. Black tiles show optimal Bartlett correction factor closest to 1 of the individual parameter in each column. Since columns are ordered by this correction factor, the x-axis corresponds to the approximative conservativeness ratio *aCR*. White line and black bold numbers indicate the uncorrected outcome for *C* = 1, whereas green line and numbers show overall optimal Bartlett correction for a *CR* ≈ 95%. Deviations from 0.95 on x-axis originate from binning issues. (A) Results for all 768 parameters, (B) same results grouped by parameter type and (C) identifiability status. (D) Outcome for individual models reveal that a comprehensive overall optimal correction factor is difficult to determine. (E) Overall optimal Bartlett correction factors *C*\* from exhaustive search with resulting thresholds 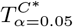 for the likelihood ratio statistic.

The heatmaps in Fig. 11 allow to determine an *overall optimal* Bartlett-like correction factor *C*\* for all analyzed parameters, but also separate overall correction factors *C*\* of the groups of the specific model parameter types, based on their identifiability status or individually for each benchmark model. This *overall optimal*correction factor *C*\* is defined such that an appropriate correction yields a conservativeness ratio *CR* = 0.95 or close to this value. This corresponds to 95% of the parameters in the heatmap classified preferably as perfect consensus case or at least conservative. The *overall optimal* Bartlett-like correction factor *C*\* can be visually determined in the heatmaps by intersecting the vertical green line at *aCR* = 0.95 or its closest value with the black curve of individual optimal correction factors. This procedure implies that all parameters on the left-hand side of the intersection are classified as *at least conservative* for the respective correction factor.

In fact, for all examples in Fig. 11, the approximative conservativeness ratio *aCR* matches the exact conservativeness ratio CR, since the non-monotonic classifications occur only for the upper (right hand sided) 5% of the columns. The green numbers on the y-axis of the heatmaps indicate the *C*\* of the respective parameter subset.

The respective statistical threshold 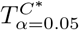 for the likelihood-ratio test can be determined from the Bartlett correction factor when assuming that the corrected likelihood ratio with the respective overall optimal correction factor, is now 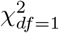-distributed, cf. Eq. (21). Fig. 11E summarizes the overall optimal correction factors for each parameter group and model with the resulting alternative statistical thresholds 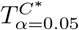 which could be used in applications instead of the typically assumed T_α=0.05_ = 3.84 from Wilks’ threorem. The analysis of the individual benchmark models shows a large variability with corrected alternative threshold values 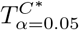 ranging from approximately 3.9 to 18.5.

It appears that a generally applicable correction factor *C*\*, i.e. for all parameter types as well as for the group of only dynamic parameters would be *C*\* = 0.52. For the parameter groups of only observational and initial value parameters are closer to the uncorrected scenario with *C*\* = 0.66 and *C*\* = 0.69, respectively. Error parameters, however, require a rather strong correction of the likelihood ratios, indicated by a Bartlett correction factor of *C*\* = 0. 21, resulting in a rather high value of the threshold 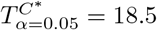.

The overall analysis analysis for all investigated parameters yields 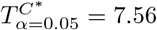 for an alternative Bartlett-corrected statistical threshold. However, the analysis of parameter groups and individual models reveals a large variability of the optimal correction factors, depending on the specific application, i.e. model and experimental design setting. Thus, despite the detailed and extensive search approach it remains difficult to identify a reliable overall Bartlett-like correction factor and threshold that is generally applicable in most situations.

#### Adapted thresholds using Chebyshev’s and Cantelli’s inequality

In order to enable a more appropriate description of the likelihood ratio statistic, another approach for an alternative approximation of their unknown distribution is employed that might be used in realistic application scenarios.

While the *empirical rule* allows for approximative boundaries for e.g. confidence intervals in the case of Gaussian or approximatively bell-shaped distributions, *Chebyshev’s* and *Cantelli’s* inequality can be used for any kind of distribution under mild conditions. Here, they are specifically used to draw valid conclusions about the statistical thresholds also in the finite-sample case. For the empirical likelihood ratios, it was shown in the previous analyses that a 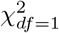-distribution in principle is a reasonable choice to describe the likelihood-ratio test statistic. Thus, the idea is to use both inequalities with an in detail unknown, but 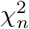-like distribution. Only the mean *μ* = *n* = 1 and the variance *σ*^2^ = 2*n* = 2 of such a 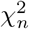-like distribution with *n* = 1 degrees of freedom need to be plugged into both inequalities for the following conclusions. This reduces the required distributional assumptions drastically without discarding the in principle appropriate 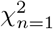-assumption entirely.

It follows from Chebyshev’s inequality in the *kσ*-neighbourhood for *k* = 4.47 around the mean that the probability of a random variable drawn from this *in detail unknown* distribution is 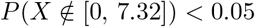. Using *k* = 4.47 this leads to an alternative threshold of 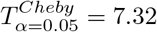 for a 5% significance level or by using *k* = 3.16 a threshold 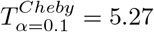 for a 10% significance level. Likewise, the typically assumed asymptotic threshold 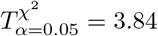 for a significance level *α* = 0.05 following Wilk’s theorem corresponds to an alternative Chebyshev-adapted threshold 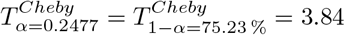 with significance level *α* = 0.2477. This corresponds to an actual confidence level of only 75.23 % instead of the desired 95 %.

In contrast to Chebyshev’s inequality, Cantelli’s inequality is restricted to a positive one-tailed distribution. It yields similar, yet slightly smaller values for the thresholds 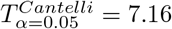 and 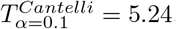. The asymptotic theory threshold value for *α* = 0.05 yields an alternative Cantelli-adapted threshold 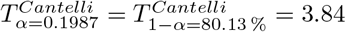 for significance level *α* = 0.1987. Again the resulting confidence level under these assumptions is only 80.13% instead of the expected confidence level of 95% from asymptotic theory.

Using these alternative thresholds, the conservativeness ratio *CR* is evaluated for the empirical likelihood ratios from the bootstrapping procedure and visualized in Fig. 12 for the previously discussed parameter groups and also for the individual models. In terms of the pp-plot it is checked whether the graph at *p_theo_* = 0.95, implying a threshold of *T* = 3.84 in the theoretical *χ*^2^-distribution corresponds to an empirical cumulative probability of at least *p_emp_* > 0.8013 = 1 – *α* for the Cantelli-adapted threshold 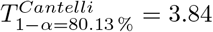. In this scenario, the empirical sample has at least *p_emp_* > 80.13% of its values below the threshold *T* = 3.84. The pp-plots are analyzed and classified as described earlier, so that a conservativeness ratio *CR* can be derived for the respective parameters. Fig. 12C illustrates the results for the threshold value *T* = 3.84 with the Cantelli interpretation of the confidence level. Likewise, it is shown how often the non-problematic classifications occur for the empirical likelihood ratios of the respective parameter groups or models for *T* = 3.84, but with the probability interpretation from asymptotic theory in Fig. 12A and and from Chebyshev’s inequality in Fig. 12 F. For larger values of the alternative thresholds, e.g. in Fig. 12 D,E,G,H the same pp-plots are evaluated at the corresponding larger *p_theo_*-values of the theoretical CDF of the 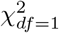-distribution and checked against the probabilities *p_emp_* = 0.9 or *p_emp_* = 0.95 of the ECDF of the empirical likelihood ratios.

**Fig 12.**
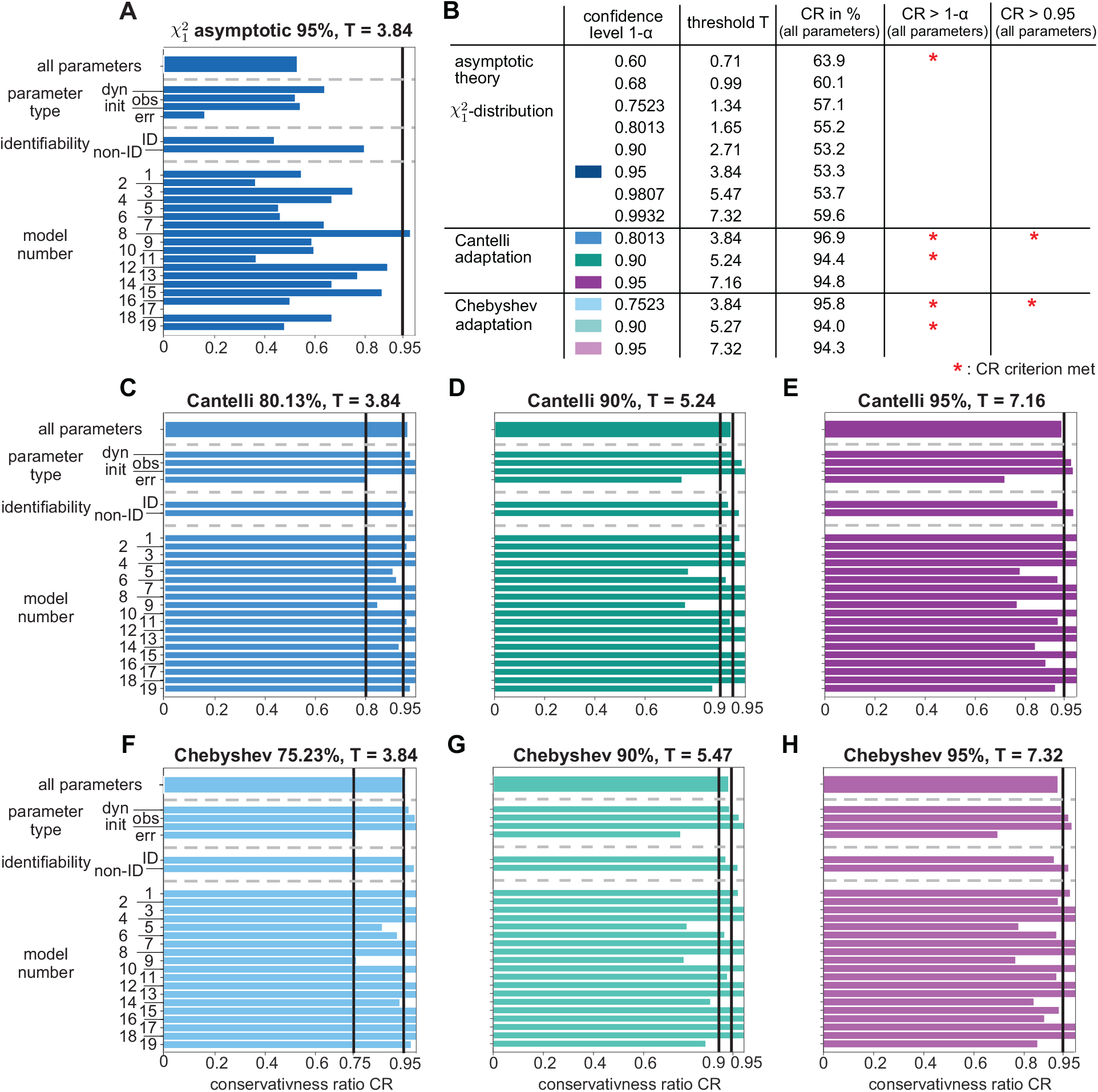
Conservativeness ratio *CR* at asymptotic and adapted thresholds. (A) Histogram of conservativeness ratio CR for the typically utilized 95%-threshold from asymptotic theory for all analyzed parameters, separately for parameter groups and individual models as a reference. (B) Table summarizes the CR for all parameters for alternative confidence levels using asymptotic theory and for adapted thresholds or adapted confidence levels using Chebyshev’s or Cantelli’s inequality. (C, F) All blue histograms including panel (A) share the same threshold value *T* = 3.84, but differ in the interpretation of the confidence level. (D, G) The green histograms correspond to a confidence level of 90% with slightly different thresholds using Chebyshev’s or Cantelli’s inequality. (E, H) Likewise, purple histograms correspond to a confidence level of 95%. Vertical black solid lines in the histograms indicate the *CR* > 1 – *α* and *CR* > 0.95 limit, which coincides for the asymptotic case.

Overall, these adapted thresholds show a preferable conservativeness ratio *CR* for most parameter groups and models. In contrast, the asymptotic theory threshold in Fig. 12A does by far not reach either the *CR* > 0.95, nor the *CR* > 1 – *α* limit, except for one single model especially error parameters are far off from a desirable *CR*. The largest confidence level for which at least *CR* > 1 – *α* holds on average for all investigated parameters using asymptotic theory is 1 – *α* = 0.6, cf. Fig. 12 B. Using the adapted thresholds, the weaker limit of *CR* > 1 – *α* is reached for both, the Chebyshev- and the Cantelli-based re-interpretation of the asymptotic threshold value *T* = 3.84 in all parameter groups and models. Both criteria are met for the overall *CR* for all parameters and for most models. Error parameters, which were often classified as problematic during this study, now show a conservativeness ratio, which is sufficient for the *CR* > 1 – *α* criterion in Fig. 12C and F. When increasing the confidence level, however, error parameters remain problematic, also when using the Chebyshev- or Cantelli-adaptation.

In general, the adapted threshold for a confidence level of 95% shows only a minor deviation in the *CR* from the required 95%-limit on average for all parameters, cf. Fig. 12 B, E and H. Thus, 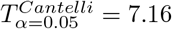 could serve as an appropriate *alternative threshold* for likelihood ratios in the finite-sample case for the 95 %-confidence level in the future. For existing studies, the typically utilized threshold value of T = 3.84 for an originally assumed confidence level of 1 – α = 95% could be re-interpreted in this line for the finite-sample case using the *alternative confidence level* 1 – *α* ≥ 80.13% from the Cantelli-adaptation.

## Conclusions

The analysis of empirical likelihood ratios from 19 nonlinear ODE benchmark models reveals a general deviation of their distribution from asymptotic theory for more than half of the estimated parameters. Thus, the usage of *Wilks’ Theorem* as an asymptotic approximation of the likelihood-ratio test might be problematic in typical ODE modeling applications with limited amounts of experimental data. In this finite-sample case, the typically utilized asymptotic thresholds of the test statistic are anti-conservative in approximately 45% of the analyzed cases. For these parameters, for example profile likelihood-based confidence intervals tend to be underestimated, i.e. the confidence intervals are too small.

The appropriateness of the asymptotic assumption depends on the actual model system and experimental setup. The results of this work show a large variability between the benchmark models and between the parameters within the models. No pattern in the model or parameter properties could be identified that indicates a preferable accordance of the asymptotic thresholds for the likelihood ratios in realistic applications. The only exception are error model parameters, which have been identified as rather problematic, i.e. they are anti-conservative in most cases and thus require special attention.

The magnitude of the deviation from the asymptotic case however is small enough in most cases so that suitable corrections can be applied in order to increase the fraction of *at least* conservative cases, when using likelihood ratio statistic-related measures. In principle, the Bartlett correction could be applied in such scenarios, but for nonlinear ODE models these correction factors are only available through a numerical search approach. The presented comprehensive analysis of all analyzed parameters, yields a candidate for an finite-sample corrected threshold for the likelihood ratios through a Bartlett correction factor for which 95% of all parameters show at least conservative results. The threshold value from this extensive search Bartlett correction is close to another alternative threshold adaptation, motivated from Cantelli’s inequality. There, the adapted threshold for the 95 %-confidence level in typical application settings has a value of 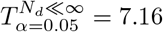 and could be used as a finite-sample alternative for the asymptotic threshold of 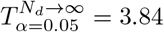 for future studies.

Such an alternative threshold formulation can be likewise utilized for a re-interpretation of results in existing studies, which employ the asymptotic threshold *T* = 3.84 for the likelihood-ratio test for example in likelihood profiles. When utilizing the asymptotic threshold value *T* = 3.84, a re-interpretation of the confidence level based on Cantelli’s inequality of approximately 80% in a typical finite-sample case seems more appropriate in contrast to the commonly assumed confidence level of 95 %.

For practical applications, three *modi operandi* in decreasing order of recommendation are plausible. It is most recommended to adopt statistical methods based on likelihood ratios and to use one of the adapted thresholds values, for example 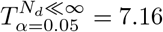 motivated from Cantelli’s inequality instead of *T*_*α*=0.05_ = 3.84, in order to avoid anti-conservative cases. Alternatively and preferable for results in existing studies, the commonly assumed asymptotic threshold 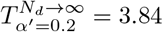 is utilized, but the results are re-interpreted by a more realistic confidence level of only 80% instead of 95%. Lastly and least favorable, nothing is changed in the usage of the asymptotic threshold value of 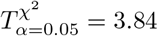 and assuming a 95% confidence level, but the presence of approximately 47% of anti-conservative cases and for example too small confidence intervals simply needs to be accepted.

## Data availability statement

All data, including model definition files and originally published parameters, all data realisations used for the bootstrapping approach, all related scripts, fits and results and all code to analyse the results and to generate the plots is available on Zenodo at https://doi.org/10.5281/zenodo.7738793.

## Author Contributions

Conceptualization: Bernhard Steiert, Clemens Kreutz.

Investigation: Christian Tönsing.

Methodology: Christian Tönsing, Bernhard Steiert, Jens Timmer, Clemens Kreutz.

Software: Christian Tönsing.

Supervision: Jens Timmer, Clemens Kreutz.

Writing - original draft: Christian Tönsing.

Writing - review & editing: Christian Tönsing, Jens Timmer, Clemens Kreutz.

## Acknowledgments

The authors acknowledge support by the state of Baden-Württemberg through bwHPC and the German Research Foundation (DFG) through grant INST 35/1134-1 FUGG. C.T and C.K. were supported by the German Research Foundation (DFG) under Germany’s Excellence Strategy - EXC-2189 - project ID: 390939984. C.K. was supported by the German Ministry of Education and Research by grant EA:Sys [FKZ031L0080].

## Supporting information

**S1 Fig. Parametric bootstrapping procedure for empirical likelihood ratios.**

### Algorithm 1 Parametric bootstrapping procedure for empirical likelihood-ratios

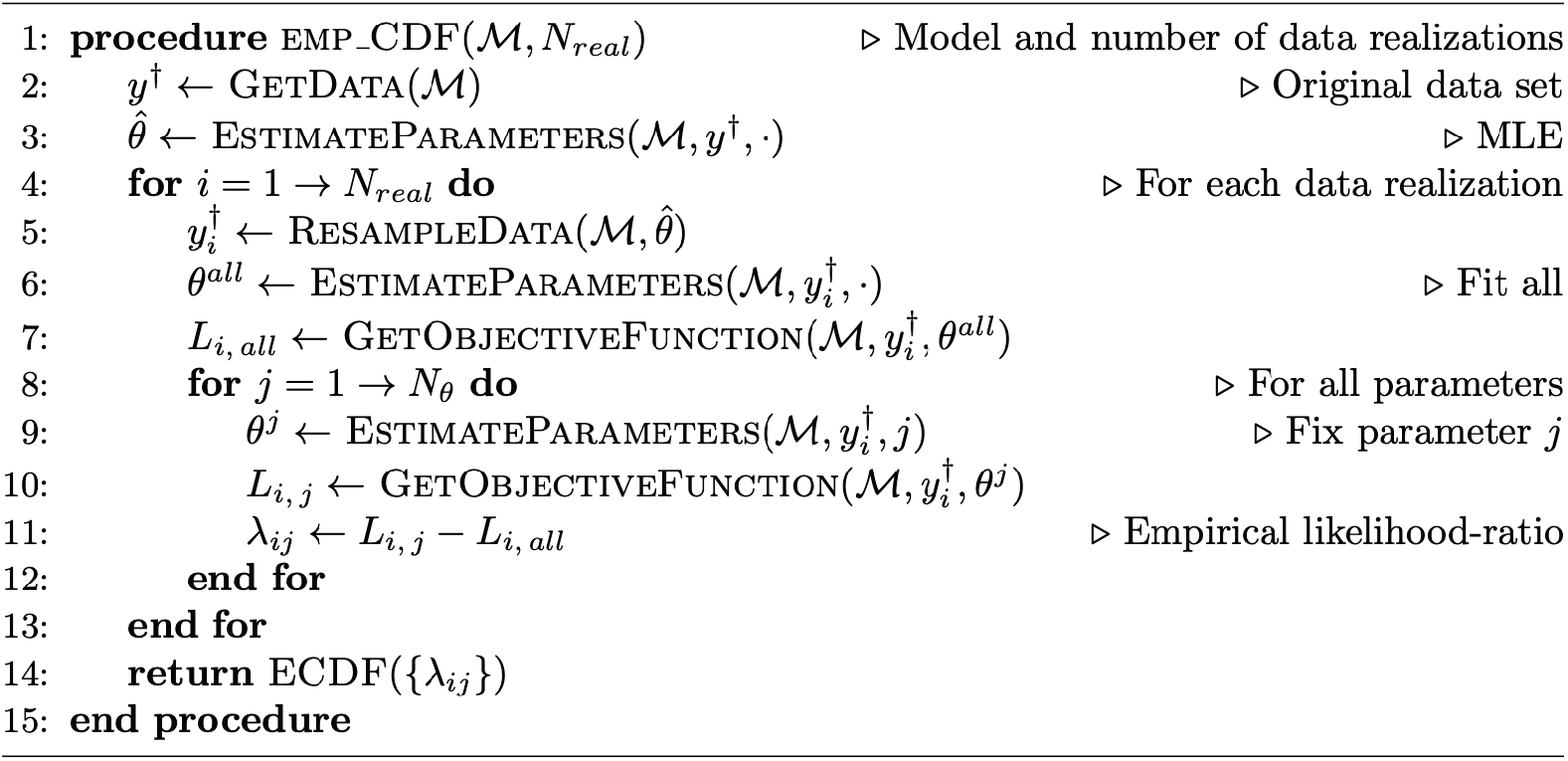

## References

1. Schoeberl B, Eichler-Jonsson C, Gilles ED, Müller G. Computational modeling of the dynamics of the MAP kinase cascade activated by surface and internalized EGF receptors. Nature Biotechnology. 2002;20(4):370.

2. Kholodenko BN. Cell-signalling dynamics in time and space. Nature Reviews Molecular Cell Biology. 2006;7(3):165–176.

3. McHugh ML. The chi-square test of independence. Biochemia medica. 2013;23(2):143–149.

4. Berger JO, Wolpert RL. The likelihood principle: A review and generalizations. Department of Statistics, Purdue University; 1982.

5. Neyman J, Pearson ES. On the problem of the most efficient tests of statistical hypotheses. Philosophical Transactions of the Royal Society of London Series A, Containing Papers of a Mathematical or Physical Character. 1933;231(694-706):289–337.

6. Kreutz C, Raue A, Timmer J. Likelihood based observability analysis and confidence intervals for predictions of dynamic models. BMC Systems Biology. 2012;6(1):120.

7. Ward EJ. A review and comparison of four commonly used Bayesian and maximum likelihood model selection tools. Ecological Modelling. 2008;211(1-2):1–10.

8. Raue A, Kreutz C, Maiwald T, Bachmann J, Schilling M, Klingmüller U, et al. Structural and practical identifiability analysis of partially observed dynamical models by exploiting the profile likelihood. Bioinformatics. 2009;25(15):1923–1929.

9. Maiwald T, Hass H, Steiert B, Vanlier J, Engesser R, Raue A, et al. Driving the model to its limit: Profile likelihood based model reduction. PLOS ONE. 2016;11(9):e0162366.

10. Tönsing C, Timmer J, Kreutz C. Profile likelihood-based analyses of infectious disease models. Statistical Methods in Medical Research. 2018;27(7):1979–1998.

11. Raue A, Schilling M, Bachmann J, Matteson A, Schelke M, Kaschek D, et al. Lessons learned from quantitative dynamical modeling in Systems Biology. PLOS ONE. 2013;8(9):e74335.

12. Kreutz C, Timmer J. Systems Biology: Experimental design. FEBS Journal. 2009;276(4):923–942.

13. Kitano H. Foundations of systems biology. The MIT Press Cambridge, Massachusetts London, England; 2001.

14. Cox DR, Hinkley DV. Theoretical Statistics. Chapman & Hall; 2000.

15. Lehmann EL, Casella G. Theory of Point Estimation. Springer Science & Business Media; 2006.

16. Strutz T. Data Fitting and Uncertainty: A Practical Introduction to Weighted Least Squares and Beyond. Vieweg and Teubner; 2010.

17. Cordeiro GM, Cribari-Neto F. An Introduction to Bartlett Correction and Bias Reduction. Springer; 2014.

18. Wilks SS. The large-sample distribution of the likelihood ratio for testing composite hypotheses. Annals of Mathematical Statistics. 1938;9(1):60–62.

19. Honerkamp J. Statistical Physics: An Advanced Approach with Applications. Springer; 2002.

20. Kreutz C, Timmer J. Optimal Experiment Design, Fisher Information. In: Encyclopedia of Systems Biology. Springer; 2013. p. 1576–1579. Available from: https://doi.org/10.1007/978-1-4419-9863-7_1222.

21. Venzon DJ, Moolgavkar SH. Method for computing profile-likelihood-based confidence intervals. Journal of the Royal Statistical Society: Series C. 1988;37(1):87–94.

22. Murphy SA, van der Vaart AW. On profile likelihood. Journal of the American Statistical Association. 2000;95(450):449–465.

23. Kreutz C, Raue A, Kaschek D, Timmer J. Profile likelihood in Systems Biology. FEBS Journal. 2013;280(11):2564–2571.

24. Raue A, Becker V, Klingmüller U, Timmer J. Identifiability and observability analysis for experimental design in nonlinear dynamical models. Chaos. 2010;20(4):045105.

25. Bartlett MS. Properties of sufficiency and statistical tests. Proceedings of the Royal Society of London Series A. 1937;160(901):268–282.

26. DiCiccio T, Hall P, Romano J, et al. Empirical likelihood is Bartlett-correctable. Annals of Statistics. 1991;19(2):1053–1061.

27. Lawley DN. A general method for approximating to the distribution of likelihood ratio criteria. Biometrika. 1956;43(3/4):295–303.

28. Cordeiro GM. Improved likelihood ratio statistics for generalized linear models. Journal of the Royal Statistical Society: Series B. 1983;45(3):404–413.

29. Bayer FM, Cribari-Neto F. Bartlett corrections in beta regression models. Journal of Statistical Planning and Inference. 2013;143(3):531–547.

30. Melo TF, Ferrari SL, Cribari-Neto F. Improved testing inference in mixed linear models. Computational Statistics & Data Analysis. 2009;53(7):2573–2582.

31. Johansen S. A Bartlett correction factor for tests on the cointegrating relations. Econometric Theory. 2000;16(5):740–778.

32. Chernick MR. The Essentials of Biostatistics for Physicians, Nurses, and Clinicians. Wiley Online Library; 2011.

33. Chebyshev PL. Des valeurs moyennes. Journal de Mathématiques Pures et Appliquées. 1867;12(2):177–184.

34. Cantelli FP. Sui confini della probabilita. In: Atti del Congresso Internazionale dei Matematici: Bologna del 3 al 10 de settembre di 1928; 1929. p. 47–60.

35. Efron B, Tibshirani RJ. An Introduction to the Bootstrap. Chapman and Hall; 1993.

36. Gibbons JD, Chakraborti S. Nonparametric Statistical Inference: Revised and Expanded. CRC Press; 2014.

37. Alkan O, Schoeberl B, Shah M, Koshkaryev A, Heinemann T, Drummond DC, et al. Modeling chemotherapy-induced stress to identify rational combination therapies in the DNA damage response pathway. Science Signaling. 2018;11(540):eaat0229.

38. Bachmann J, Raue A, Schilling M, Böhm ME, Kreutz C, Kaschek D, et al. Division of labor by dual feedback regulators controls JAK2/STAT5 signaling over broad ligand range. Molecular Systems Biology. 2011;7:516.

39. Becker V, Schilling M, Bachmann J, Baumann U, Raue A, Maiwald T, et al. Covering a broad dynamic range: information processing at the erythropoietin receptor. Science. 2010;328(5984):1404–1408.

40. Boehm ME, Adlung L, Schilling M, Roth S, Klingmüller U, Lehmann WD. Identification of isoform-specific dynamics in phosphorylation-dependent STAT5 dimerization by quantitative mass spectrometry and mathematical modeling. Journal of Proteome Research. 2014;13(12):5685–5694.

41. Brännmark C, Palmer R, Glad ST, Cedersund G, Strålfors P. Mass and information feedbacks through receptor endocytosis govern insulin signaling as revealed using a parameter-free modeling framework. Journal of Biological Chemistry. 2010;285(26):20171–20179.

42. Bruno M, Koschmieder J, Wuest F, Schaub P, Fehling-Kaschek M, Timmer J, et al. Enzymatic study on AtCCD4 and AtCCD7 and their potential to form acyclic regulatory metabolites. Journal of Experimental Botany. 2016;67(21):5993–6005.

43. Fiedler A, Raeth S, Theis FJ, Hausser A, Hasenauer J. Tailored parameter optimization methods for ordinary differential equation models with steady-state constraints. BMC Systems Biology. 2016;10(1):80.

44. Hass H, Kipkeew F, Gauhar A, Bouché E, May P, Timmer J, et al. Mathematical model of early Reelin-induced Src family kinase-mediated signaling. PLOS ONE. 2017;12(10):e0186927.

45. Isensee J, Kaufholz M, Knape MJ, Hasenauer J, Hammerich H, Gonczarowska-Jorge H, et al. PKA-RII subunit phosphorylation precedes activation by cAMP and regulates activity termination. Journal of Cell Biology. 2018;217(6):2167–2184.

46. Lucarelli P, Schilling M, Kreutz C, Vlasov A, Boehm ME, Iwamoto N, et al. Resolving the combinatorial complexity of Smad protein complex formation and its link to gene expression. Cell Systems. 2018;6(1):75–89.

47. Merkle R, Steiert B, Salopiata F, Depner S, Raue A, Iwamoto N, et al. Identification of cell type-specific differences in erythropoietin receptor signaling in primary erythroid and lung cancer cells. PLOS Computational Biology. 2016;12(8):e1005049.

48. Kurzhunov D, Borowiak R, Hass H, Wagner P, Krafft AJ, Timmer J, et al. Quantification of oxygen metabolic rates in Human brain with dynamic ^17^O MRI: Profile likelihood analysis. Magnetic Resonance in Medicine. 2017;78(3):1157–1167.

49. Raia V, Schilling M, Böhm M, Hahn B, Kowarsch A, Raue A, et al. Dynamic Mathematical Modeling of IL13-Induced Signaling in Hodgkin and Primary Mediastinal B-Cell Lymphoma Allows Prediction of Therapeutic Targets. Cancer Research. 2011;71(3):693–704.

50. Schwen LO, Schenk A, Kreutz C, Timmer J, Rodriguez MMB, Kuepfer L, et al. Representative sinusoids for hepatic four-scale pharmacokinetics simulations. PLOS ONE. 2015;10(7):e0133653.

51. Sneyd J, Dufour JF. A dynamic model of the type-2 inositol trisphosphate receptor. Proceedings of the National Academy of Sciences. 2002;99(4):2398–2403.

52. Swameye I, Müller T, Timmer J, Sandra O, Klingmüller U. Identification of nucleocytoplasmic cycling as a remote sensor in cellular signaling by databased modeling. Proceedings of the National Academy of Sciences. 2003;100(3):1028–1033.

53. Zheng Y, Sweet SM, Popovic R, Martinez-Garcia E, Tipton JD, Thomas PM, et al. Total kinetic analysis reveals how combinatorial methylation patterns are established on lysines 27 and 36 of histone H3. Proceedings of the National Academy of Sciences. 2012;109(34):13549–13554.

54. Hass H, Loos C, Raimúndez-Álvarez E, Timmer J, Hasenauer J, Kreutz C. Benchmark problems for dynamic modeling of intracellular processes. Bioinformatics. 2019;35(17):3073–3082.

55. Raue A, Steiert B, Schelker M, Kreutz C, Maiwald T, Hass H, et al. Data2Dynamics: a modeling environment tailored to parameter estimation in dynamical systems. Bioinformatics. 2015;31(21):3558–3560.

56. Xie M, Singh K. Confidence distribution, the frequentist distribution estimator of a parameter: A review. International Statistical Review. 2013;81(1):3–39.

57. Bjerhammar A. Application of Calculus of Matrices to Method of Least Squares: With Special Reference to Geodetic Calculations. Elander Göteborg; 1951.

58. Transtrum MK, Machta BB, Sethna JP. Geometry of nonlinear least squares with applications to sloppy models and optimization. Physical Review E. 2011;83(3):036701.

59. Lill D, Timmer J, Kaschek D. Local Riemannian geometry of model manifolds and its implications for practical parameter identifiability. PLOS ONE. 2019;14(6).

60. Drton M. Likelihood ratio tests and singularities. Annals of Statistics. 2009;37(2):979–1012.

61. Steiert B, Raue A, Timmer J, Kreutz C. Experimental Design for Parameter Estimation of Gene Regulatory Networks. PLOS ONE. 2012;7(7):e40052.

62. Tönsing C, Timmer J, Kreutz C. Cause and cure of sloppiness in ordinary differential equation models. Physical Review E. 2014;90(2):023303.

